# Efficient generation of lower induced Motor Neurons by coupling *Ngn2* expression with developmental cues

**DOI:** 10.1101/2022.01.12.476020

**Authors:** Francesco Limone, Jana M. Mitchell, Irune Guerra San Juan, Janell L. M. Smith, Kavya Raghunathan, Alexander Couto, Sulagna Dia Ghosh, Daniel Meyer, Curtis J. Mello, James Nemesh, Brittany M. Smith, Steven McCarroll, Olli Pietiläinen, Ralda Nehme, Kevin Eggan

## Abstract

Human pluripotent stem cells (hPSCs) are a powerful tool for disease modelling and drug discovery, especially when access to primary tissue is limited, such as in the brain. Current neuronal differentiation approaches use either small molecules for directed differentiation or transcription-factor-mediated programming. In this study we coupled the overexpression of the neuralising transcription factor Neurogenin2 (*Ngn2*) with small molecule patterning to differentiate hPSCs into lower induced Motor Neurons (liMoNes). We showed that this approach induced activation of the motor neuron (MN) specific transcription factor *Hb9/MNX1*, using an*Hb9*::GFP-reporter line, with up to 95% of cells becoming *Hb9*::GFP^+^. These cells acquired and maintained expression of canonical early and mature MN markers. Molecular and functional profiling revealed that liMoNes resembled bona fide hPSC-derived MN differentiated by conventional small molecule patterning. liMoNes exhibited spontaneous electrical activity, expressed synaptic markers and formed contacts with muscle cells *in vitro*. Pooled, multiplex single-cell RNA sequencing on 50 cell lines revealed multiple anatomically distinct MN subtypes of cervical and brachial, limb-innervating MNs in reproducible quantities. We conclude that combining small molecule patterning with Ngn2 can facilitate the high-yield, robust and reproducible production of multiple disease-relevant MN subtypes, which is fundamental in the path to propel forward our knowledge of motoneuron biology and its disruption in disease.

## INTRODUCTION

Since their discovery, many groups have recognised the prodigious abilities of stem cells to differentiate into almost any cell type of the body. This unique differentiation capability, or pluripotency, can facilitate the understanding of basic biology and malfunctions of tissues that are hard to access and that are specifically highly evolved in humans, such as the Central Nervous System (CNS) (Penney et al., 2020). Most neuronal differentiation schemes mimic developmental embryonic signals by small molecule patterning. The neuralisation of stem cells is achieved by manipulating bone morphogenic protein (BMP) and transforming growth factor *β* (TGF*β*), commonly referred to as “dual-Smad inhibition” (Chambers et al., 2009). This initial study further showed that different combinations of small molecules used as patterning factors could push neuronal progenitors towards distinct neuronal fates. From there, many have developed and refined differentiation protocols for specific neuronal subtypes. However, caveats still remain, such as: the incomplete neuralisation of cultures, underlining the need for additional neuralising factors (Chambers et al., 2012); the particularly long time needed to generate fully mature neuronal cultures and the heterogeneity in differentiation efficiency between cell lines (Boulting et al., 2011; Kampmann, 2020).

To overcome these limitations, others have employed different approaches which couple stem cell technologies with the overexpression of a neuralising transcription factors (TFs) (Mertens et al., 2016). These TFs have been used to generate induced Neurons (iNs) from fibroblasts (Vierbuchen et al., 2010), and the combination with subtype-specific TFs was able to generate specific types of neurons (Son et al., 2011). These approaches have been applied and translated to stem cells with one of the more recent reports of Ngn2 being able to differentiate human Pluripotent Stem Cells (hPSCs) into glutamatergic neurons (Zhang et al., 2013). These advances allowed reproducible generation of neurons in shorter time and in fewer steps. However, these reprogramming methods may skip several of the pivotal developmental steps that are part of normal neuronal specification. For this reason, questions have been raised regarding the actual heterogeneity of the identity of the generated cell populations and of the impact of the enforced overexpression of TFs to downstream applications (Lin et al., 2021).

We wanted to generate spinal Motor Neurons (MNs) for biological modelling of degenerative motoneuron diseases, such as Amyotrophic Lateral Sclerosis (ALS) and Spinal Muscular Atrophy (SMA) that selectively affect these highly specialised neuron types (Han et al., 2011). MNs are the only neurons to exit the nervous system and contact skeletal muscles to allow us to breathe and move through a specific synaptic contact, the Neuro-Muscular Junction (NMJ). Protocols to differentiate this neuronal subtype are based on decades of developmental biology studies (Jessell, 2000; Stifani, 2014) and are extensively reviewed elsewhere (Davis-Dusenbery et al., 2014; Sances et al., 2016). These protocols entail neuralization inputs described above coupled with ventralising factors like Sonic Hedgehog and/or its agonists (Shh/SAG) and the caudalising effects of retinoids (retinoic acid – RA) (Amoroso et al., 2013; Boulting et al., 2011; Dimos et al., 2008; Kiskinis et al., 2014) or, alternatively, by overexpressing a combination of transcription factors: *Ngn2, Isl1, Lhx3* (also known as NILs) (De Santis et al., 2018; Hester et al., 2011). Both approaches have proven to be useful for investigations in MN biology. However, on one hand directed differentiation produces cultures containing different cell types other than MNs with high line-to-line heterogeneity rendering disease modelling studies difficult. On the other hand, the overexpression of three TFs produces pure cultures but very specific subtypes of MNs. This limits the scalability of these studies since several, specific combinations of TFs are needed to reproduce the diversity of MN subtypes *in vitro*.

Previously, we have demonstrated that overexpression of Ngn2 coupled by small molecules patterning is able to enhance the regional specification of neurons from hPSCs to the cortical-like patterned induced Neurons - *piNs* (Nehme et al., 2018). Additionally, small molecules have also been reported to enhance efficiency of MN programming (Hester et al., 2011; Mazzoni et al., 2013b). This led us to hypothesize that the coupling of Ngn2 expression alone with ventralising SAG and caudalising RA could efficiently generate MN-like cells. Here, we report that the addition of different patterning molecules during Ngn2-programming of hPSCs can lead to specification of regionally defined neuronal states. With time in culture, differentially patterned cells developed into morphologically different neurons that maintain regionally defined transcriptomic features according to small molecules treatment. A reporter cell line for the MN-specific transcription factor *MNX1/Hb9* demonstrated that ∼95% of the cells subjected to SAG and RA activated this master regulator TF for MN development. This, in combination with the expression of pan-MN markers validated the cellular identity of SAG- and RA-patterned-Ngn2 cells as MN-like cells: the lower induced Motor Neurons (liMoNes). liMoNes not only expressed canonical markers of MN and resembled bona fide hiPSC-derived MNs, they were also electrophysiologically active and able to form synaptic contact with muscle cells *in vitro*. By leveraging newly developed analysis tools for single-cell RNA-sequencing (scRNAseq) technology that allow analysis of dozens cell lines cultured in the same dish at the same time, we demonstrated that the protocol can spontaneously produce several subtypes of disease-relevant facial and limb-innervating MNs in an exceptionally robust fashion, reproducible across 47 stem cell lines. This combinatorial approach addressed several shortcomings found in previously published protocols and will facilitate the use of these cells in the understanding of basic spinal cord MNs biology and its disruption in disease.

## RESULTS

### Ngn2-driven neuralization can be directed to different neuronal fates by small molecules patterning

Given that the combination of patterning molecules with neuralising Ngn2 expression could generate cortical excitatory neurons (Nehme et al., 2018), we wondered whether the protocol could be repurposed with the use of alternative patterning factors, such as RA and SAG, to generate other types of neurons, i.e. MNs. To test this hypothesis, we set out to optimise an Ngn2-based protocol that would substitute WNT inhibition, used to generate cortical neurons (piNs) (ibidem), with ventralising SAG and caudalising RA to induce a ventral-posterior fate and potentially generate lower-induced Motor Neurons (liMoNes) (Figures 1A-B).

**Figure 1.**
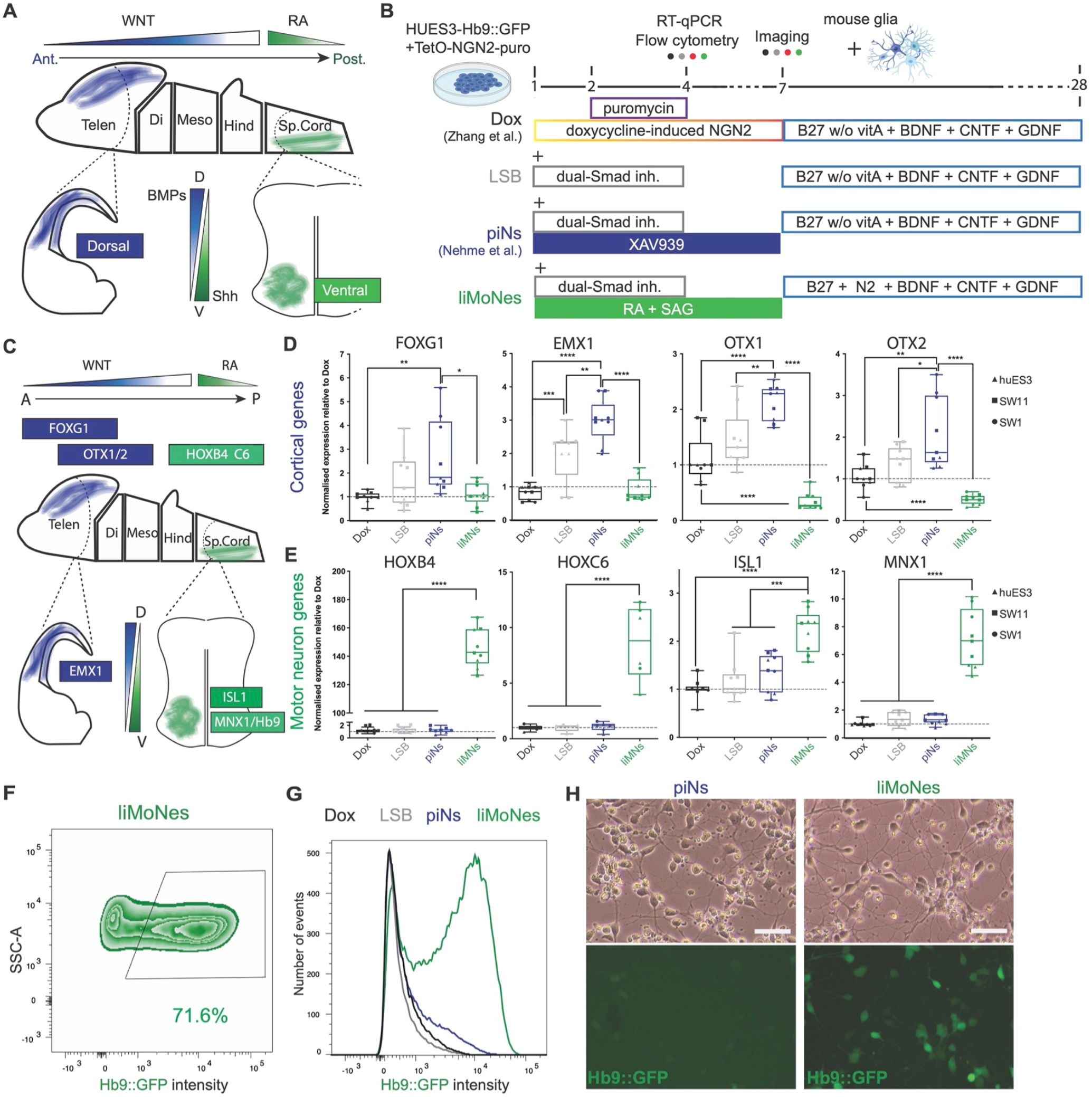
Ngn2-driven neuralization can be directed to different neuronal fates by small molecules patterning. (A) Diagram of known developmental cues used to design patterning strategy. (B) Differentiation schemes used for comparison of divergent Ngn2-driven trajectories: Dax - original Ngn2 overexpression from Zhang et al. 2013; LSB - Ngn2 overexpression coupled with neuralising dual-Smad inhibition (LDN193189, S6431542); piNs - cortical-like patterned induced Neurons (Nehme et al. 2018); liMoNes - lower induced Motor Neurons generated by Ngn2-overexpression and ventro-caudal patterning (Retinoic Acid and Smoothened Agonist). (C) genes selected as master regulators of anterior-dorsal, cortical development and ventro-caudal, spinal cord development. (D) RT-qPCR quantification for induction of cortical genes after rostro-dorsalising WNT inhibition (p-values from one-way ANOVA). (E) RT-qPCR quantification demonstrating induction of spinal genes after caudal-ventralising induction of SAG and RA (one-way ANOVA). (F) Flow cytometry quantification of Hb9::GFP positive cells at day 4. (G) Hb9::GFP intensity at day4 of differentiation demonstrating higher total intensity of the Hb9::GFP signal in liMNs (H) Hb9::GFP expression day7 post-induction in piNs and liMoNes, the majority of liMoNes express the reporter (scale bar 50 micron).

To test if the patterning induced regionally specified neuronal states, we selected TFs pivotal for early neuronal development that are specifically divergent between the cortex and the spinal cord (Figures 1C). To this end we collected RNA early into the protocol and performed RT-qPCR, at a stage described as Neuronal Progenitor Cell (NPC)-like (Nehme et al., 2018), to assess the expression of these TFs. While rostro-dorsalising WNT inhibition induced the expression of master regulators of cortical development *EMX1*, *FOXG1*, *OTX1* and *OTX2* (Figure 1D), the caudal-ventral patterning induced the expression of posterior markers *HOXB4* and *HOXC6*, of the cholinergic master regulator *ISL1* (Pfaff et al., 1996) and specifically of *MNX1* (Hb9), expressed only by spinal motor neurons in the nervous system (Arber et al., 1999) (Figure 1E). Importantly, caudal-ventral patterning significantly reduced the expression of *OTX1* and *OTX2*, transcription factors that regulate the schism between the cortex and more posterior regions of the CNS (Muhr et al., 1999). In line with previous studies, dual-Smad inhibition in combination with Ngn2 was followed by loss of transcription factors associated with pluripotency, *OCT4* and *SOX2*, and acquisition of early neuronal markers, *PAX6* and *TUBB3* (Figures S1A-B) (Nehme et al., 2018).

To further confirm the regional specification of these NPCs, we took advantage of a human embryonic stem cell line engineered with a GFP reporter under the MN-specific Hb9 promoter (Di Giorgio et al., 2007). Flow cytometry analysis of GFP^+^ cells confirmed that by day 4 after doxycycline-induced Ngn2 overexpression, more than 70% of cells treated with RA and SAG were GFP positive (Figures 1F). Strikingly, not only was the percentage of GFP^+^ cells higher, but the intensity of GFP signal increased with ventro-caudal patterning (Figure 1G), in agreement with the higher levels of *MNX1*/Hb9 RNA. By day 7, cells subjected to RA and SAG showed strong *Hb9::GFP* expression whereas only a few, very dim GFP positive cells were visible in the other conditions (Figures 1H and S1C-D). Taken together, the data from early neuronal stages suggest that differential patterning coupled with Ngn2-overexpression leads to the specification of different neuronal progenitor fates, including the MN fate, *in vitro*.

### The neuronal fate induced by patterned Ngn2 expression is maintained throughout differentiation

We then proceeded to confirm that the regional specification was maintained long-term during the differentiation. For this purpose, we extended *in vitro* culturing by replating cells in neuronally supportive conditions (Figure 2A). First, we analysed cell morphology by microscopy. Patterning produced neurons with strikingly different morphology with piNs showing small, polarised cell bodies and MN-patterned cells showing a wider, multipolar shape with one extended axon (Figure 2B and S2A-B), strikingly reminiscent of the morphology of cortical pyramidal neurons and spinal, ventral-horn motor neurons *in vivo*, respectively (Abdel-Maguid and Bowsher, 1984).

**Figure 2.**
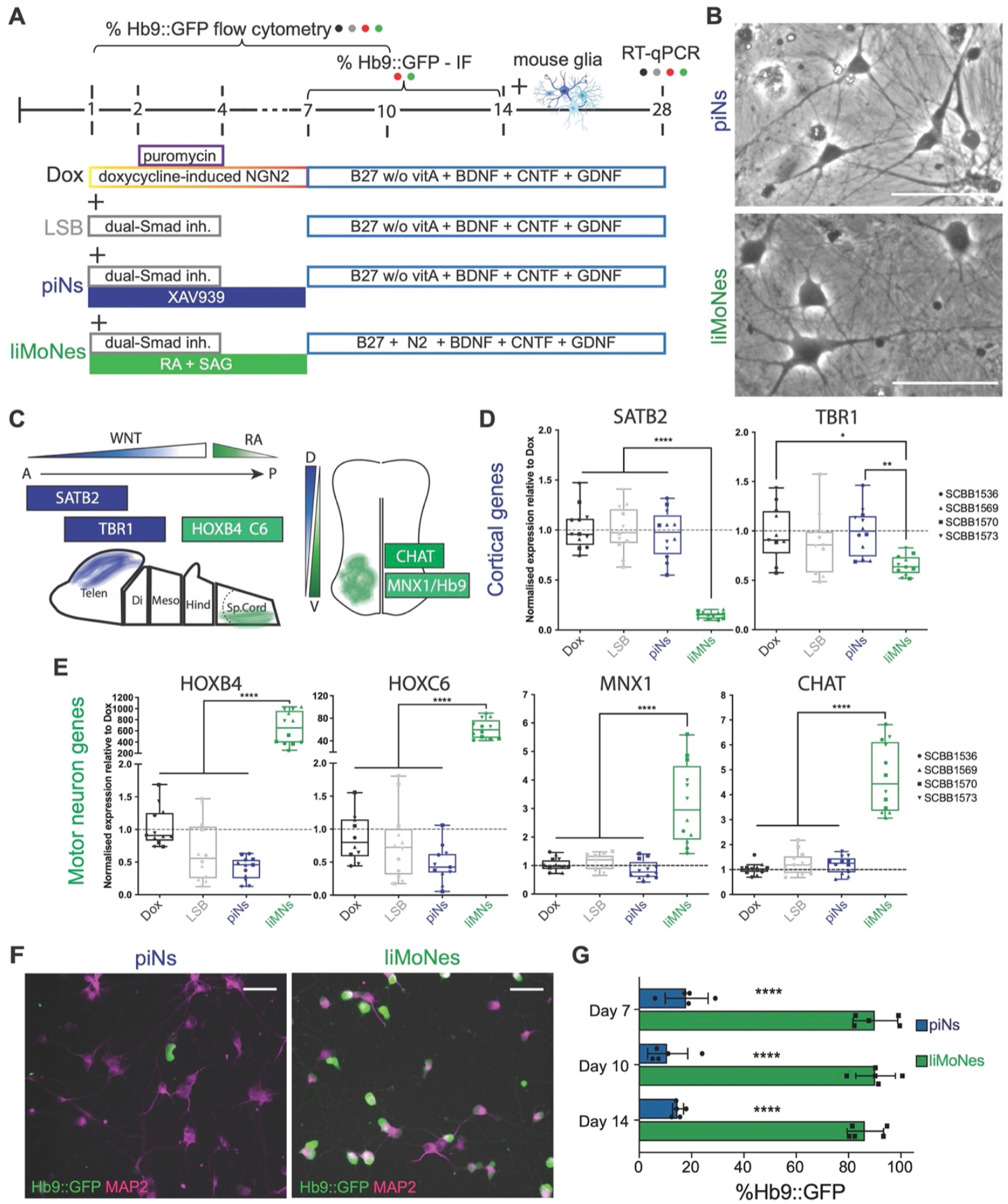
Patterned Ngn2-induced neuronal fate is maintained throughout the differentiation. (A) Differentiation schemes implemented for neuronal maturation following one-week-long patterning: Dox - original Zhang et al. 2013; LSB - Ngn2 with dual-Smad inhibition; piNs - cortical-like patterned induced Neurons (Nehme et al. 2018); liMoNes - lower induced Motor Neurons. (B) Imaging at day30 of piNs and liMoNes (scale bar 100 micron). (C) diagram of genes specifically only expressed in either anterior-dorsal cortical neurons or ventro-caudal, spinal cord motor neurons. (D) RT-qPCR quantification for induction of cortical genes in piNs (p-values from one-way ANOVA). (E) RT-qPCR quantification demonstrating induction of cortical genes in liMoNes one-way ANOVA). (F) Hb9::GFP expression at day14 post-induction in piNs and liMoNes (scale bar 50 micron). (G) Quantification of Hb9::GFP expression at day7, 10 and 14 post-induction in piNs (blue) and liMoNes (green) by immunofluorescence (p-values from I-test at each time point).

To confirm that the regional identity specified by patterning was maintained, we collected RNA at day 30 of differentiation and investigated the expression of genes known to be specifically expressed in either glutamatergic neurons of the cortex or cholinergic motor neurons of the spinal cord (Figure 2C). We confirmed that the caudalisation resulted in repression of the cortical genes *SATB2* and *TBR1* (Figure 2D). Expression of posterior markers *HOXB4* and *HOXC6* was sustained in caudalised cells and slightly suppressed in piNs (Figure 2E). Moreover, more mature ventralised cells expressed the MN-specific transcription factor, *MNX1* (*Hb9*) and higher transcript levels of the main component of cholinergic machinery, Choline Acetyltransferase (*CHAT*) (Figure 2E), while maintaining expression of panneuronal markers (Figure S2C). According to this polarised gene expression, expression of the *Hb9::GFP* reporter was also maintained through-out the differentiation only in RA- and SAG-patterned cells, reaching a peak of ~95% at day 7 (Figure 2F-G and S2D-E), and was then slightly downregulated as seen in early development of MNs of the spinal cord *in vivo* (Thaler et al., 2004). The data so far confirmed that, as per the NPC-like stage, coupling of Ngn2 overexpression with patterning factors can produce regionally specified neurons and we define the ventralised and caudalised cultures as lower-induced Motor Neurons: *liMoNes*.

### liMoNes reproducibly express canonical pan-Motor Neuron markers and resemble bona fide hPSC-MN

Given that neuralisation by Ngn2 overexpression can be directed to different neuronal fates and maintained during *in vitro* culture, we wanted to confirm the expression of key motor neuron markers at the protein level. By day 30, liMoNes expressed Cholinergic Acetyltransferase (Chat) (Figure 3A) and limb-innervating specific marker Foxp1 (Figure 3B) (Dasen et al., 2008). Moreover, liMoNes showed reactivity for antibodies against the transcription factor Islet1 along with SMI-32, that recognises MN-enriched neurofilament heavy chain (Figure 3C). These two, together with Hb9, form the trifecta recognised as the human pan-Motor Neuron staining (Amoroso et al., 2013). Indeed, 60-90% of cells express at least one these markers (figure 3D), while the pan-MN markers were robustly and reproducibly expressed by 80-90% of cells by different cell lines (Figure 3E).

**Figure 3.**
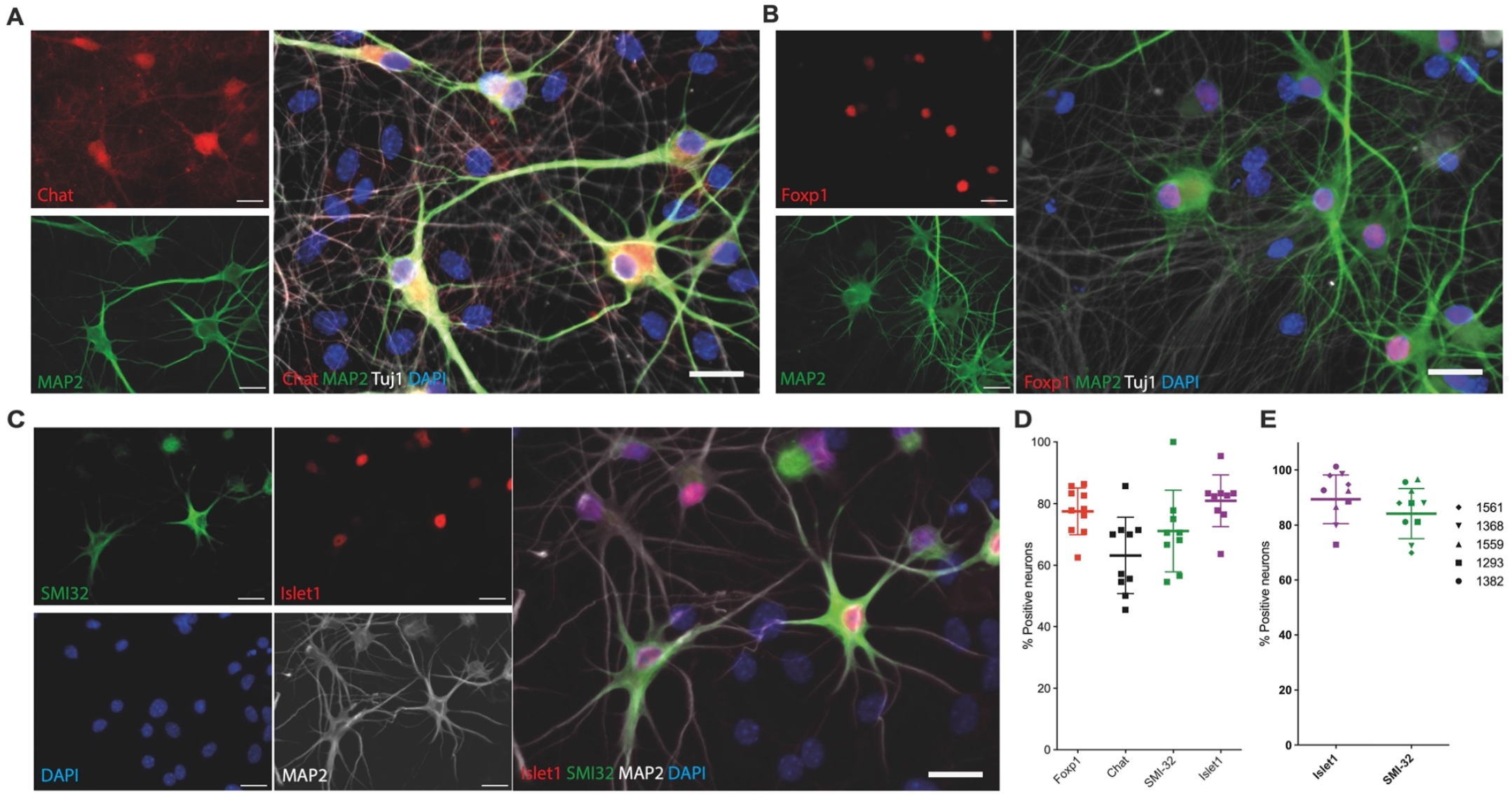
liMoNes reproducibly express canonical Motor Neuron markers. (A) lmmunofluorescence analysis for cholinergic marker Chat and neuronal cytoskeletal proteins MAP2 and Tubb3 (Tuj1) (scale bar 30 micron). (B) lmmunofluorescence analysis for Limb-innervating MN marker Foxp1 and neuronal MAP2 and Tubb3 (Tuj1) (scale bar 30 micron). (C) lmmunofluorescence analysis for MN-enriched SMl-32, cholinergic transcription factor lslet1 and neuronal MAP2 (scale bar 30 micron). (D) Quantification for cells in A-C. (E) Quantification of expression of selected markers in five independently differentiated lines.

We next wanted to confirm that liMoNes resembled cells defined by the scientific community as bona fide hiPSC-derived Motor Neurons. We thus differentiated MNs following a conventional, widely used method using neuralising small molecules and patterning factors (*2D MN*) (Klim et al., 2019). Briefly, stem cells were subjected to neuralising dual-Smad inhibition followed by DAPT and SU5402 while caudalised and ventralised with RA and SAG. Differentiated neurons were separated from the mixed cultures by sorting for cell surface marker N-cam 14 days post-neuronal induction (Klim et al., 2019), and then cultured in neuronal differentiation media, under similar conditions to liMoNes for 14 more days (Figure 4A). We then compared the morphologies of the conventional 2D MNs and liMoNes by imaging. We found that morphology of liMoNes was similar to that of 2D MN, with large multipolar cell bodies which was distinct from the morphology of cortical cells (Figure 4B). Moreover, liMoNes and 2D MN expressed similar patterns of pan-MN staining (Figure S3A-B). Remarkably, RT-qPCR analysis revealed that liMoNes expressed even higher transcript levels of limb-innervating motor neurons marker *HOXC6*, and comparable levels of other motor-neuron markers (Figure 4C). These results confirmed that liMoNes resemble one kind of bona fide hiPSC-derived motor neurons defined by the broader scientific community.

**Figure 4.**
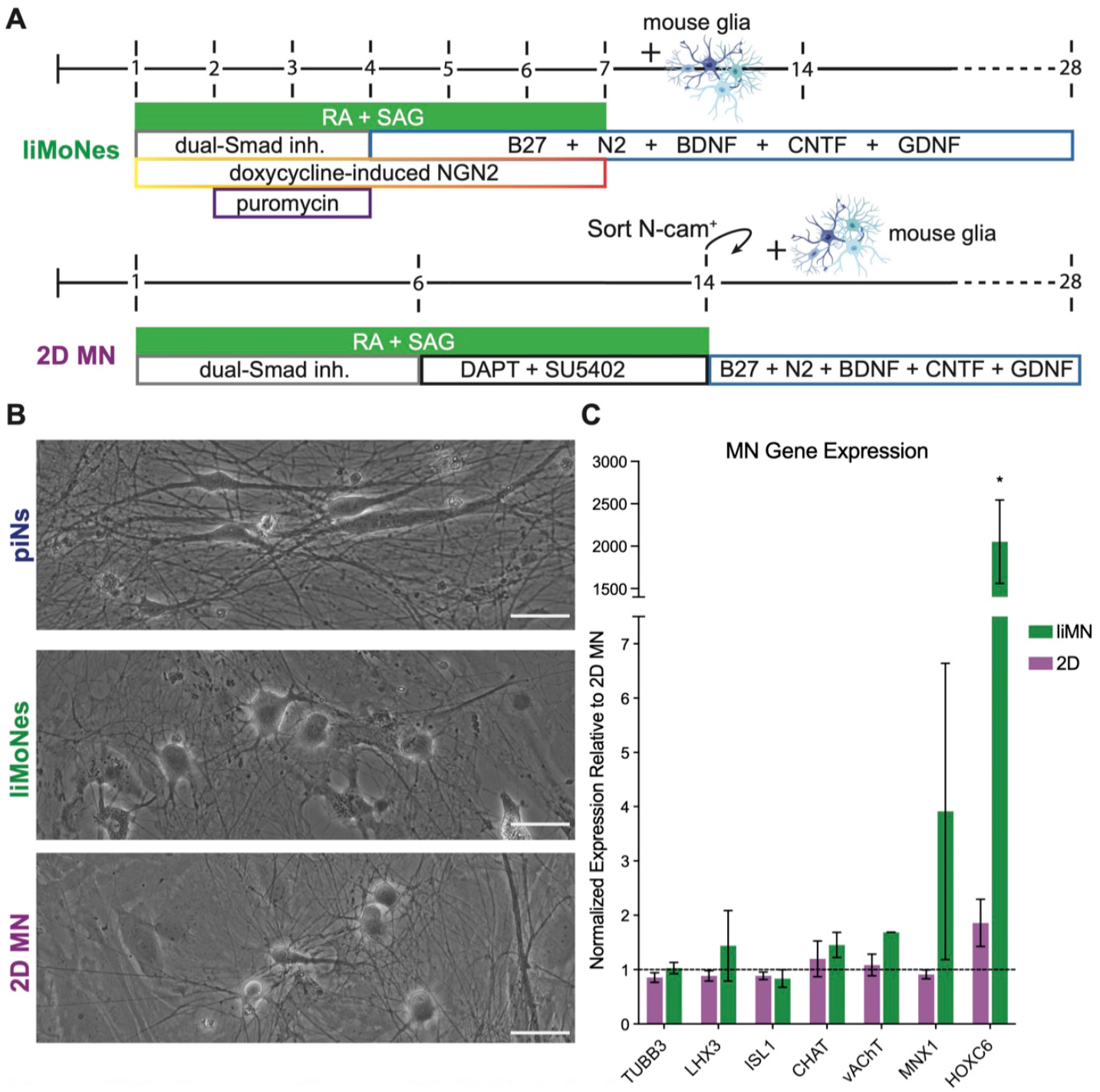
liMoNes resemble bona fide hPSC-derived motor neurons. (A) Differentiation schemes implemented to compare liMoNes with bona fide MN derived from pluripotent cells by conventional small molecule induction (2D MN, in purple). (B) Morphology of neuronal cells produced: piNs, liMoNes and 2D-MN (scale bar 50 micron). (C) RT-qPCR quantification of MN markers between liMoNes (green) and 2D-MN (purple).

### liMoNes form active synaptic networks and contact muscle cells*in vitro*

To investigate the functional properties of liMoNes, we set out to assess their ability to form synapses. By day 30 of culture, liMoNes displayed abundant punctate staining for the basic presynaptic component Synapsin along neurites and one neurite positive for Axonal-marker AnkyrinG (Figure 5A). This cultures also showed protein expression of other pre- and post-synaptic molecules, denoting the formation of an architecturally sound synaptic network similar to the one previously described in piNs (Figure S4A-B, (Nehme et al., 2018)). We then used multielectrode arrays (MEAs) to assess their electrophysiological properties. The neuronal cultures showed a steady increase in spiking rates over time (Figure S4C-D). Moreover, treating cells with potassium-gated channel opener Retigabine, a regulator of electrophysiology of MNs and a therapeutic agent for Amyotrophic Lateral Sclerosis (Wainger et al., 2014; Wainger et al., 2021), efficiently silenced the cultures underlining the usefulness of these cells for the study of neurodegenerative diseases (Figure 5B).

**Figure 5.**
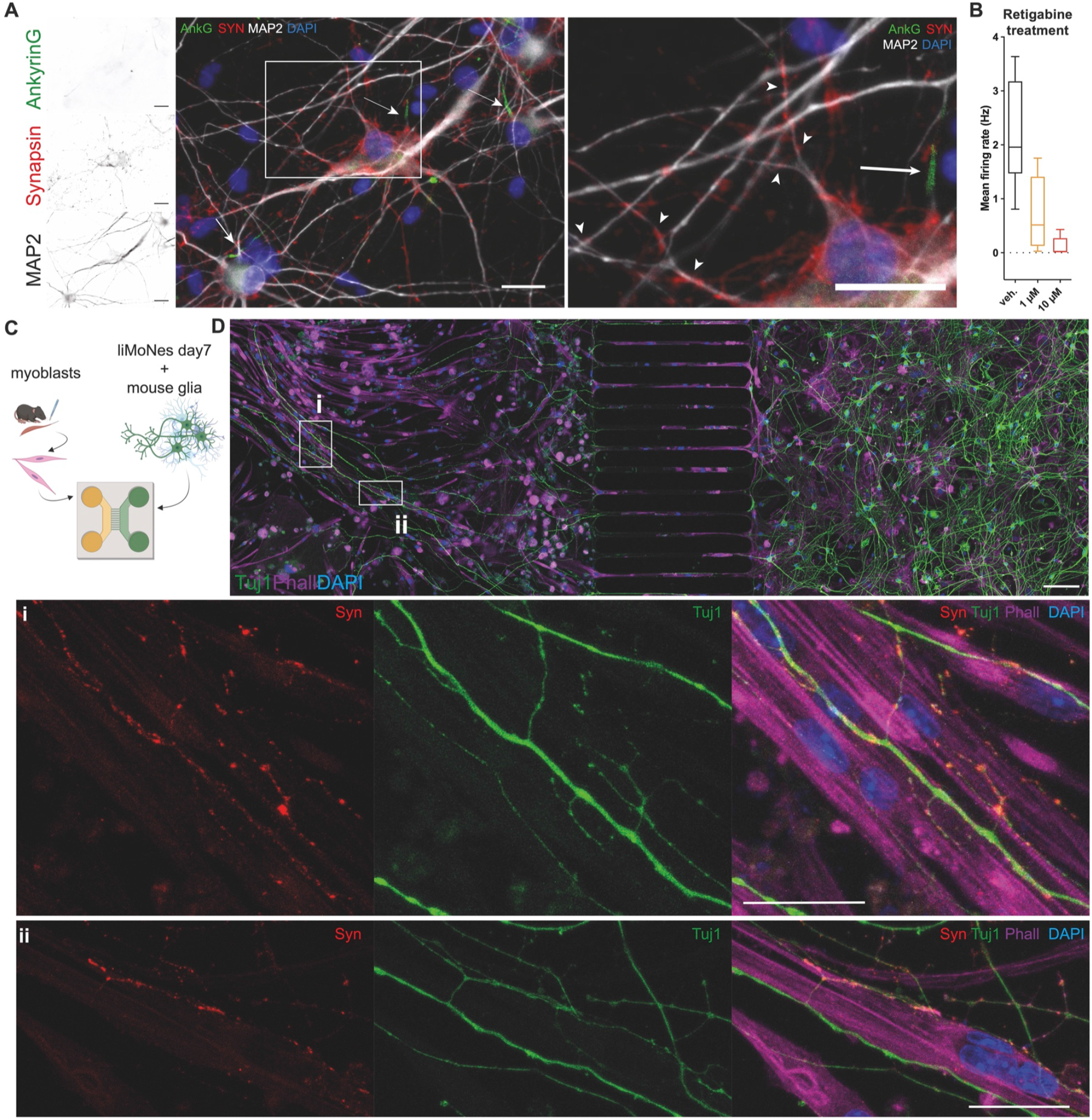
liMoNes can form active synaptic structures in vitro. (A) liMoNes express proteins involved in the formation of functional axons and synaptic structure. (B) Mean number of spikes in day50 cultures treated with raising concentrations of Retigabine. (C) Diagram of co-culture experiments of liMoNes and primary murine myoblasts in microfluidic devices. (D) lmmunofluorescence of co-culture of liMoNes and primary murine myoblasts showing glia-liMoNes co-cultures (right), where neurons extend axons through the channels (middle), contacting primary muscle cells (left). (D^i^-D^ii^) Insets of (D) showing liMoNes forming synaptic-like contacts with muscles cells (scale bar 50 micron).

Motor neurons are very specialised cells being the only neurons to exit the nervous system and contact muscles through a highly specific synaptic contact: the NMJ. To test the ability of liMoNes to form NMJ-like structure we established co-cultures with murine muscle cells in compartmentalised microfluidic devices where neurons grown in one chamber are free to extend axons through groves that connect them to muscle cells grown in the second chambers (Figure 5C). Staining of these cultures showed that liMoNes successfully extended axonal projections on the other side of the device and contact muscle (Figure 5D and S5A). Strikingly, these projections can form structures reminiscent of the NMJ shape along muscle cells that expressed pre-synaptic protein Synapsin (Figure 5D^i^-D^ii^ and S5B-E), a sign of first development of synaptic contact, even though they did not achieve clustering of acetylcholine receptors on muscle cells. So far, only one other study has shown this clustering is possible *in vitro* using culture conditions that recapitulate the physical separation between MN cell bodies and muscle (Stoklund Dittlau et al., 2021) and only with manipulations of medium/gradients of molecules and prolonged time in culture.

### scRNA-seq confirms expression of MN-specific genes and reproducibility of the protocol

After confirming the MN-like properties of liMoNes, we set out to characterise their molecular identity and reproducibility by single cell RNA sequencing. We coupled sequencing with two newly developed technologies: *Census-seq* and *Dropulation* (Mitchell et al., 2020; Wells et al., 2021) to enable the characterization of dozens biological replicates in a single experiment. These methods utilise the intrinsic variability of single nucleotide polymorphisms (SNPs) within a population as a barcode to assign identities in a mixed culture - a “village” - of multiple donors, similarly to pooled CRISPR-Cas9 barcoded screens (Shalem et al., 2014; Wang et al., 2014). More precisely, *Census-seq* allows population-scale, quantitative identity assignment from a mixed group of donors (Mitchell et al., 2020), *Dropulation* can assign identities at a single cell level in a “village” for RNA sequencing studies (Wells et al., 2021). With this aim in mind, we produced liMoNes “villages”: 50 embryonic stem cell lines, previously subjected to whole-genome sequencing, were separately differentiated into liMoNes. At day 7 post-induction, postmitotic cells were pooled in equal numbers to make up “villages” containing all donors in one dish. Using genotypes from WGS we were able to reassign the donor identities in a mixed village (Figure 6A).

**Figure 6.**
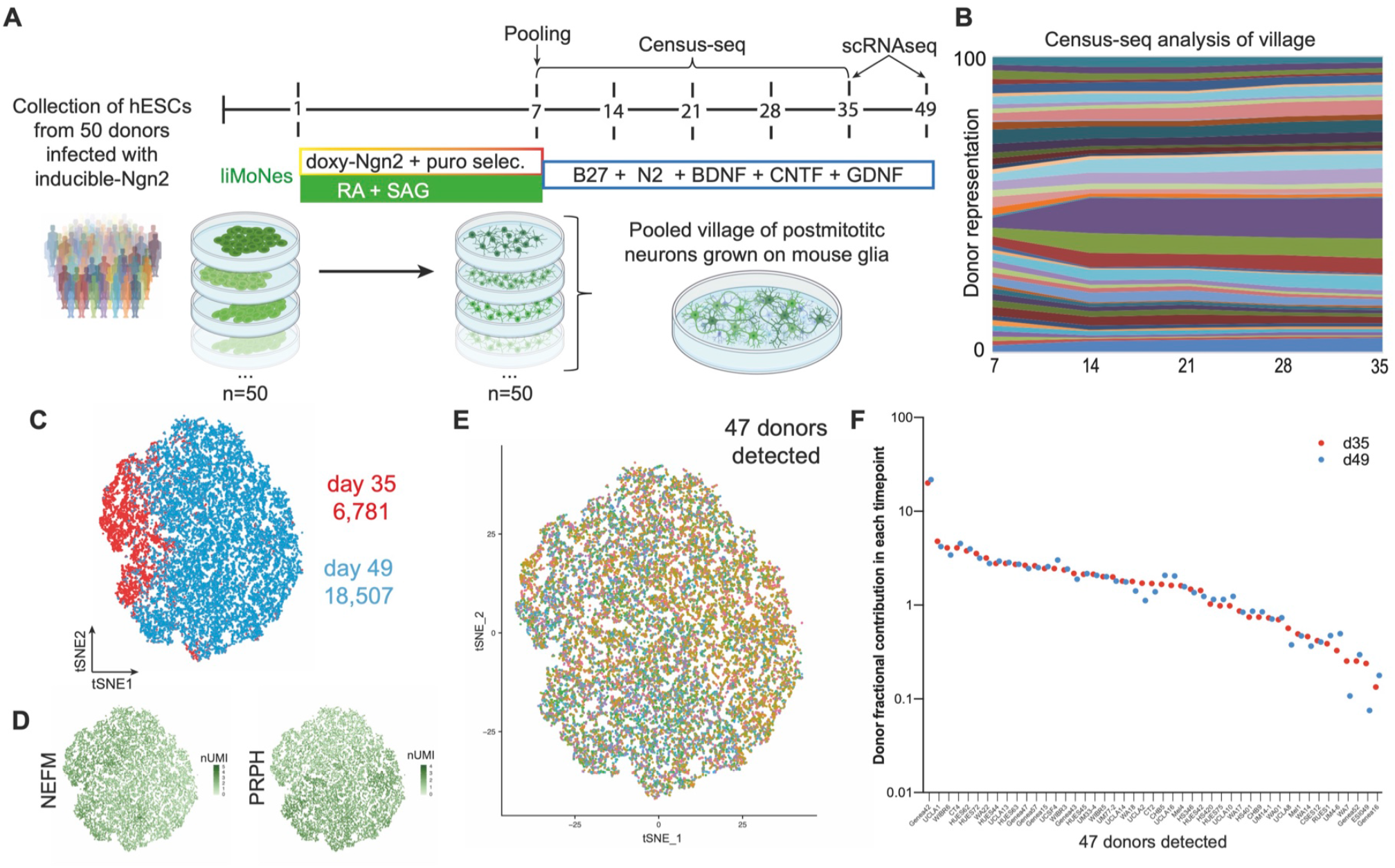
scRNA-seq confirms expression of MN-specific genes and reproducibility of the protocol. (A) Pooling strategy and village construction for *Census-seq* and *Dropulation* analysis. (B) Sandplot of *Census-seq* analysis showing balanced representation of47 detected donors throughout several days post-induction. (C) *t*-SNE projection of scRNAseq analysis of 25,288 cells of two timepoints of mature liMoNes differentiation. (D) *t*-SNE projection with expression of markers for neurons of the peripheral nervous system. (E) *t*-SNE projection of 25,288 cells depicting donor’s identity of each cell from47 donors detected by *Dropulation* analysis. (F) Fraction representation of47 donors in the two timepoints of mature liMoNes differentiation.

To ensure that the donor composition remained balanced throughout the differentiation, cells were harvested once a week to collect genomic DNA for low-coverage sequencing. *Census-seq* analyses showed that we could detect 47 of the 50 donors originally pooled and confirmed that donor distribution remained consistent for at least four weeks (Figure 6B). Neurons were then harvested at day 35 and day 49 for scRNA-seq and *Dropulation* analysis. Libraries generated from 25,288 cells demonstrated strong expression of neuronal markers, especially of the peripheral nervous system (PNS), *NEFM* and *PRPH* (Figure 6C-D and S6A). Using the newly devised *Dropulation* analytical pipeline, we were able to assign donor identity to barcoded droplets. Initial *t*-SNE clustering showed an even distribution of each donor (Figure 6E) and we confirmed that the contribution of each donor to the villages remained constant at both timepoints (Figure 6F) underlying the robustness and reproducibility of the protocol.

### Ventro-caudal patterning of Ngn2 can produce different MN subtypes

After establishing that the protocol can generate a similar culture of cells robustly and reproducibly in 47 cell lines, we studied the transcriptomic identities of liMoNes. Firstly, liMoNes expressed typical MN markers including *STMN2*, *NEFH*, *ISL1* and *MNX1* (Figure S6B). Next, we focused on genes indicative of cholinergic identity and found low but detectable expression of *ACHE*, *SLC5A*7 (Cht1), *SLC18A3* (vAChT) (Figure S6C). Finally, we detected expression of *AGRN* and *NRG1*, genes expressed by MNs and involved in the formation of NMJs (Figure S6D). We therefore confirmed that our protocol can reproducibly generate MN-like cells from all 47 cell lines.

*In vivo*, motor neurons are usually classified in different anatomical subtypes (a.k.a. pools or columns) according to their position along the spinal cord and the anatomical part of the body they connect to. Four main groups lie in parts of the spinal cord whose development is regulated by retinoids: 1. Medial Motor Column (MMC) located along the entire spinal cord and connected to axial musculature to maintain body position, 2. Spinal Accessory Column (SAC) in the cervical area innervates head and neck, 3. Phrenic Motor Column (PMC), also cervical, innervates the diaphragm, 4. Lateral Motor Column (LMC), in the brachial level at the cervical-thoracic boundary, connects to forelimbs and is itself divided in ventral-innervating, medial or dorsal-innervating, lateral LMC (Figure 7A) (Stifani, 2014). We studied the expression of previously identified markers of these MN subtypes to investigate whether subpopulations of liMoNes resembled distinct MN pools of the spinal cord, strikingly we were able to find several of these markers expressed in our scRNA-seq dataset: a small group of *PHOX2A* and *PHOX2B*-expressing SAC-like cells (Figure 7B), wide expression of PMC-enriched *ALCAM* and *POU3F1* (*SCIP*) (Figure 7C) and of markers of both lateral and medial LMCs: *FOXP1* and *LHX1* (Figure 7D-E).

**Figure 7.**
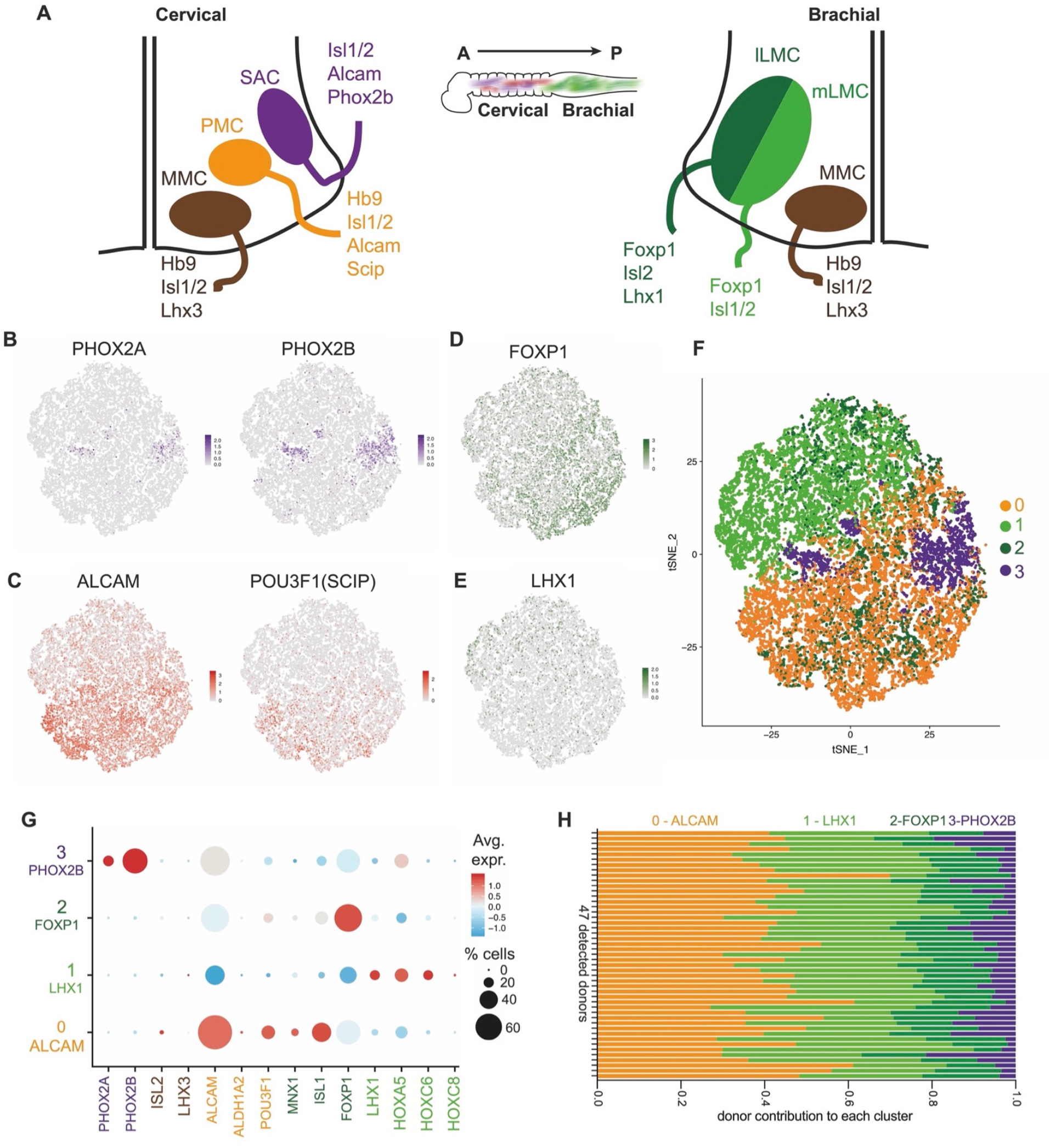
Ventro-caudal patterning of NGN2 can produce different MN subtypes. (A) Diagram of known pools of MN subtypes along mammalian spinal cord. (B) *t*-SNE projection with expression of markers specific for SAC subtypes. (C) *t*-SNE projection with expression of markers specific for PMC subtypes. (D) *t*-SNE projection with expression of markers specific for mLMC subtypes. (E) *t*-SNE projection with expression of markers specific for ILMC subtypes. (F) *t*-SNE projection of four, unbiasedly identified subclusters in the 25,288 cells analysed. (G) Dotplot for differential gene expression of MN subtype-specific markers in the four cervico-brachial MN groups. (H) Fraction of each donor’s share between the identified subclusters as calculated by *Dropulation*.

We wondered if the discrete expression of these markers would shape subgroups of cells with transcriptomic profiles that corresponded to different MN-pools. We decided to unbiasedly identify subclusters of cells and found four separate groups: liMoNes 0,1,2,3 (Figure 7F). Intriguingly, main markers for motor neuron pools segregated within the four groups and demarcated an ALCAM^+^-PMC-like group, an LHX1^+^ and a FOXP1^+^ mLMC and lLMC-like groups, and a small PHOX2B^+^ SAC-like group (Figure 7G and S7A-C). No demarcated expression for markers of the MMC pool was found (Figure S7D) consistent with reports identifying this subpopulation to be less responsive to certain patterning factors (Amoroso et al., 2013). Given the single-cell resolution of our approach, we observed two additional, striking features: 1. the expression of markers specific to anterior digit-innervating MNs *FIGN* and *CPNE3* (Figure S7E) (Mendelsohn et al., 2017); 2. the expression of a coordinated set of *HOX* genes that are activated in response to retinoids in the spinal cord (Philippidou and Dasen, 2013) and that have been identified as being specifically expressed in motor neurons (Dasen et al., 2005) (Figure S7F-G).

Taking advantage of the *Dropulation* technology, we investigated the distribution of donors in the pool within each subcluster and to our surprise each of the 47 detected donors distributed evenly within clusters highlighting the robustness and reproducibility of the protocol (Figure 7H). From this analysis, we conclude that liMoNes are composed of a plethora of motor neuron subtypes that intrinsically recapitulates pools and columns identified in the cervical and brachial spinal cord and that these subtypes can be robustly generated in a myriad of cells lines.

## DISCUSSION

In this study we describe a rapid and efficient protocol to generate human MN-like cells from hPSCs by combining the overexpression of neuralising factor Ngn2 (Zhang et al., 2013) and ventralising and caudalising small molecules patterning (Amoroso et al., 2013). We showed that different patterning molecules can direct the Ngn2-driven neuralisation into the specification of distinct neuronal fates that are maintained during *in vitro* culture. In particular, we show that ventral-caudal patterning alone induces expression of the MN-specific TF *MNX1*/Hb9 in ~90% of differentiated cells bypassing the previously used sorting methods to isolate MN from mixed cultures. The ventro-caudalised cells expressed pan-MN markers as identified *in vivo* and resembled bona fide hPSC-derived motor neurons giving them a lower motor neuron identity - hence lower induced Motor Neuron (liMoNes). liMoNes were able to generate electrophysiologically active cultures capable to form immature synaptic contact with muscle cells *in vitro*. By combining newly developed single-cell RNA-sequencing analyses tools, we demonstrated that this protocol could successfully generate a previously reported hard-to-produce neuronal cell type by a straightforward one-step programming. Additionally, the differentiation scheme is highly scalable and reproducible across 47 cell lines, and generated cultures containing a diverse population of disease-relevant MN subtypes.

The protocol here described was able to overcome some of the main issues reported in previously published differentiation schemes based on small molecules patterning. Specifically, we showed how with a single step induction, we were able to generate in only seven days, a pure population of post-mitotic neurons in which virtually all cells expressed the MN-specific marker *MNX1*/Hb9, whereas most protocols reported at least two weeks of differentiation to achieve partial expression of these reporter (Sances et al., 2016). Moreover, we demonstrated how the enforced expression of a transcription factor can achieve complete neuralisation of progenitor cells and avoid the generation of other cell types heterogeneously generated on a cell-line-to-cell-line dependent manner (Boulting et al., 2011) and how this method could be replicated in dozens of pluripotent lines. This single-step 7 day induction protocol would allow the generation of defined motor neuron cultures for *in vitro* modelling studies and avoid time-consuming and expensive cell-sorting step that was previously necessary to select relevant cell types from mixed ventral-caudal populations (Klim et al., 2019). Intriguingly, very few reports were able to establish cultures of human MNs that form NMJ-like structures *in vitro* (Demestre et al., 2015; Guo et al., 2011; Lin et al., 2019; Puttonen et al., 2015; Rigamonti et al., 2016; Umbach et al., 2012) and they achieved that by creating mixed cultures of MNs and muscle cells. So far only one report has established and characterised a co-culture system that allows NMJ formation *in vitro* using microfluidic devices which allow culture conditions more similar to human physiology, where neuronal cell bodies are secluded in the spinal cord and extend axons out of the PNS to the muscle (Stoklund Dittlau et al., 2021). The combination of our highly pure, reproducible, accelerated protocol to generate human MNs and this newly reported *in vitro* NMJ system could allow further understanding of MN-muscle connection in a physiologically relevant, human context.

Our study is one of the first reports to underline the malleability of Ngn2-based reprogramming and its ability to be directed to differential states by small molecules patterning mimicking embryonic development. We have thoroughly demonstrated in a previous report that patterning can direct Ngn2 towards a specific cortical-like state (Nehme et al., 2018), but this is the first side-by-side, systematic comparison of the ability of this programming method to diverge into different neuronal fates. Others have reported that overexpression of Ngn2 alone is able to produce an admixture of different neuronal subtypes of both the Central and Peripheral Nervous System (Lin et al., 2021), confirming that Ngn2-driven neuralisation is capable to intrinsically produce several neuronal subtypes. Here, we expand on this interesting biology showing that small molecule patterning can direct the multipotent neuralising ability of Ngn2 to populations of regionally specified neurons in a robust, reproducible manner.

Many molecular studies investigating the developmental biology of the spinal cord have shown how retinoids can specifically act as chromatin modulators and open chromatin domains in neural progenitor cells consistent with spinal cord identity and aid posteriorisation in NIL-based MN differentiation systems (Mazzoni et al., 2013b). Moreover, transcriptomic and epigenomic studies along NIL-based MN differentiation have shown that Ngn2 acts independently of the Isl1-Lhx3 heterodimers, upregulating neuralising factors that in turn open sites of chromatin that allow further specification into MN-fates (Velasco et al., 2017). Intriguingly, others have reported that overexpression of Ngn2 in fibroblasts coupled with patterning factors could generate small but relevant populations of cholinergic neurons, hinting at the malleability of this system (Liu et al., 2013). We speculate that the addition of patterning molecules to Ngn2-programming permits the opening of chromatin at a sites of MN development usually achieved by the overexpression of other TFs forming a similar epigenetic and transcriptomic landscape that allows specification into motor neuron identity. Other groups have reported that in multiple TF-based differentiation systems, addition of RA can result in the upregulation of sets of genes that the TFs alone could not achieve (Nickolls et al., 2020), confirming that a combinatorial approach might aid specification into desired cell types.

The use of only one transcription factor combined with small quantities of inexpensive patterning molecules renders this protocol more amenable to large-scale, high-throughput studies compared to previous protocols (Hester et al., 2011). The combinatorial use of multiple TFs often induces the generation of extremely specified subtypes of MNs (De Santis et al., 2018; Mazzoni et al., 2013a). And even though pure and well-defined, pool-specific combinations limit the ability of hPSC to differentiate into the intrinsic admixture of MN pools generated by retinoids/Shh and only elicits the transcriptomic programs of restricted pools of MNs (Mazzoni et al., 2013a). Even though an even more specified neuronal subpopulation might be ideal for certain studies, the need for the generation of multiple viruses expressed in a 1:1 ratio and/or bigger size viral constructs makes the use of these protocols for high-throughput, large-scale studies challenging from a financial and logistical point of view. Moreover, others have demonstrated how combinations of multiple transcription factors might take longer time to develop hPSC into neurons when compared to Ngn2-induction alone and that the timing of overexpression could interfere with the specific subtypes of neurons generated (Nickolls et al., 2020). Here we propose that a short pulse of Ngn2 overexpression coupled with patterning molecules not only reduces the number of TF needed to direct the specification of the neuralisation process but also allows intrinsic developmental processes to take place and generate the variegated diversity of MN subtypes seen in spinal cord development. Finally, given the differential susceptibility of subtypes of MNs to degenerate in certain diseases like ALS (Nijssen et al., 2017), having both resistant and susceptible populations reproducibly generated in one dish could help to further understand the dynamic process of neurodegeneration.

## ACKNOWLEDGMENTS

We would like to thank members of the Eggan’s and McCarroll’s groups for insights and useful discussions and Lee Rubin for support in the final steps of this project. We are grateful to the Harvard Stem Cell and Regenerative Biology Department (HSCRB) for the use of shared equipment and FACS facility. We are also grateful to the Stanley Center at the Broad Institute of Harvard and MIT for support with bioinformatics and cloud space. We thank BioRender for useful diagrams and images used in figures throughout this publication.

## AUTHOR CONTRIBUTION

Conception and study design F.L., O.P., R.N., K.E. Data analysis and interpretation F.L., O.P., R.N. Manuscript writing F.L., O.P., R.N., I.G.S.J. “Village” experiment construction and design, multi-cell line experiments and cell culture J.M., J.L.M.S., K.R. Immunofluorescence analysis and co-culture studies F.L., I.G.S.J., A.C., B.M.S. Conventional motor neuron experiments F.L., B.M.S., I.G.S.J. Bioinformatics analysis and scRNA-seq F.L., S.M., O.P., S.D.G., D.M., C.J.M., J.M.

## INCLUSION AND DIVERSITY

We worked to ensure diversity in experimental samples through the selection of the cell lines. We worked to ensure diversity in experimental samples through the selection of the genomic datasets. One or more of the authors of this paper self-identifies as an underrepresented ethnic minority in science. One or more of the authors of this paper self-identifies as a member of the LGBTQ+ community.

## RESOURCE AND DATA AVAILABILITY

Cell lines and reagents used in this study are deposited at the Stanley Centre for Psychiatric Research, Broad Institute of MIT and Harvard, please contact Ralda Nehme for enquiries. Matrices used for RNA sequencing analyses are available on request and further details on codes and algorithms can be found in other publications (Mitchell et al., 2020; Wells et al., 2021).

## METHOD DETAILS

### NGN2-based differentiations

Stem cells were grown in mTeSR1 (Stem Cell Technologies, 05850) and grown on Matrigel (Corning) coated pates at 37°C and 5% CO2. hPSCs were infected with TetO-Ngn2-Puro, TetO-GFP and rtTA lentiviral constructs (Zhang et al., 2013) produced by Alstem in mTeSR medium with 1 μM RoCK inhibitor Y-27632 for 24 hours. hPSs were then passaged and differentiation was started when cells reached 70-80% confluency. For the first four days of differentiation cells were grown in induction medium: DMEM/F12 (Life Technologies, 11320-033), N2 supplement (0.5%v/v, Gibco), 1X GlutaMAX (Gibco), 0.1mM Non-essential amino acid (Gibco), 0,5% glucose, doxycycline hyclate (2 μg/mL). Small molecules added: day 1 – DOX: none; LSB: 10 μM SB431543 (Custom Synthesis), 200 nM LDN193189 (Custom Synthesis); piNs: 10 μM SB431543 (Custom Synthesis), 200 nM LDN193189 (Custom Synthesis), 4 μM XAV939 (Stemgent, 04-00046); liMoNes: 10 μM SB431543 (Custom Synthesis), 200 nM LDN193189 (Custom Synthesis), 2 μM retinoic acid (Sigma) and 2 μM Smoothened agonist (Custom Synthesis). Day 2 to 4 - DOX: puromycin (5 μg/mL); LSB: puromycin (5 μg/mL), 10 μM SB431543 (Custom Synthesis), 100 nM LDN193189 (Custom Synthesis); piNs: puromycin (5 μg/mL), 10 μM SB431543 (Custom Synthesis), 100 nM LDN193189 (Custom Synthesis), 2 μM XAV939 (Stemgent, 04-00046); liMoNes: puromycin (5 μg/mL), 10 μM SB431543 (Custom Synthesis), 100 nM LDN193189 (Custom Synthesis), 1 μM retinoic acid (Sigma) and 1 μM Smoothened agonist (Custom Synthesis). On day 4 cells were dissociated using Accutase (Gibco) and replated in a 1:2 dilution to ensure puromycin selection of uninfected cells. For day 4 to 7, DOX, LSB and piNs cells were grown in neuronally supportive medium supplemented with small molecules as described above: Neurobasal (Life Technologies 21103049) supplemented with B27 supplement w/o vitA (2%v/v, Gibco), 1X GlutaMAX (Gibco), 0.1mM Non-essential amino acid (Gibco), 0,5% glucose with the addition of 10 ng/ml of BDNF, CNTF and GDNF (R&D Systems). For day 7 to 10, liMoNes were grown with small molecules as described above in neuronally supportive medium: Neurobasal (Life Technologies 21103049) supplemented with B27 supplement (2%v/v, Gibco), N2 supplement (0.5%v/v, Gibco), 1X GlutaMAX (Gibco), 0.1mM Non-essential amino acid (Gibco), 0,5% glucose with the addition of 10 ng/ml of BDNF, CNTF and GDNF (R&D Systems). On day 7, cells were dissociated using accutase and replated on glial co-cultures as described previously (Di Giorgio et al., 2007) in medium described above. From this time onwards, half-media change was performed every 2-3 days in neuronally supportive media described above with the only addition of 10 ng/ml of BDNF, CNTF and GDNF (R&D Systems). For most experiments, neurons were co-cultured with murine glial cells (50,000 cells/cm^2^) derived from postnatal brains (P0-2) as previously described (Di Giorgio et al., 2007), neurons were mixed with glia when replating day 7 cells 30,00 cells/cm^2^

### 2D MN differentiation

Stem cells were grown in mTeSR1 (Stem Cell Technologies, 05850) and grown on Matrigel (Corning) coated pates at 37°C and 5% CO2. Stem cells were differentiated to bona fide 2D Motor Neurons as previously described (Klim et al., 2019). This protocol based on the principle of neuralization by dual-Smad inhibition followed by the inhibition of NOTCH/FGF pathway both under the patterning capability of retinoids and Sonic Hedgehog. Briefly, once 90-95% confluent, stem cell medium was switched to differentiation medium: 1:1 mix of Neurobasal (Life Technologies 21103049) and DMEM/F12 (Life Technologies, 11320-033) supplemented with B27 supplement (2%v/v, Gibco), N2 supplement (0.5%v/v, Gibco), 1X GlutaMAX (Gibco), 0.1mM Non-essential amino acid (Gibco). For the first six days, differentiation medium was supplemented with 10 μM SB431543 (Custom Synthesis), 100nM LDN193189 (Custom Synthesis), 1 μM retinoic acid (Sigma) and 1 μM Smoothened agonist (Custom Synthesis). For the second week, differentiation medium was supplemented with: 5 μM DAPT (Custom Synthesis), 4 μM SU-5402 (Custom Synthesis), 1 μM retinoic acid (Sigma) and 1 μM Smoothened agonist (Custom Synthesis). To isolate neurons from mixed cultures we utilised an immune-panning based method previously described (Klim et al., 2019). At day 14, monolayers were dissociated with Accutase (Gibco) for 1 hour at 37°C. After gentle, repeated pipetting, cells were collected, spun down and resuspended in sorting buffer and filtered. Single cell suspensions were incubated with antibody against NCAM (BD Bioscience, 557919, 1:200) for 25 minutes, washed and NCAM^+^ cells were sorted with an BD FACS Aria II cell sorter. Sorted 2D MN were then plated on mouse glial cultures in motor neuron medium (Neurobasal (Life Technologies 21103049) supplemented with B27 supplement (2%v/v, Gibco), N2 supplement (0.5%v/v, Gibco), 1X GlutaMAX (Gibco), 0.1mM Non-essential amino acid (Gibco), 0,5% glucose) with the addition of 10 ng/ml of BDNF, CNTF and GDNF (R&D Systems). For most experiments, neurons were co-cultured with murine glial cells (150,000 cells/cm^2^) derived from postnatal brains (P0-2) as previously described (DI GIORGIO).

### Co-culture of Ngn2 motor neurons and mouse myoblasts in microfluidic devices

Mouse myoblasts from hindlimb skeletal muscles of young adult mice and mouse glia form neonatal mouse brains were isolated and cultured as previously described (Di Giorgio et al., 2007; Shahini et al., 2018). Microfluidic device chips (XC450, XONA Microfluidics) were designated a motor neuron compartment and a muscle compartment. The motor neuron compartment was coated with 0.1 mg/ml poly-L-ornithine (Sigma-Aldrich) in 50 mM Borate buffer, pH = 8.5 and 5 μg/ml laminin (Invitrogen), while the muscle compartment was coated with Matrigel (Corning). Day 7 Ngn2 motor neurons and mouse glia were seeded at a concentration of 100,000 neurons-200,000 glia/device. Myoblasts were seeded at a concentration of 150,000 device. Motor neurons were seeded in the motor neuron media described above with the addition of 10 ng/ml of BDNF, CNTF and GDNF (R&D Systems). For seeding and culturing the first 2 days, myoblasts were maintained in Myoblast media (DMEM/F12, 20% Foetal Bovine Serum and 10% heat-inactivated Horse Serum, and 10 ng/ml bFGF), after that, differentiation was initiated by adding myoblast differentiation media (DMEM high glucose, 5% heat-inactivated Horse Serum). Myoblast were sustained in differentiation medium for 3 days and then switched to motor neuron medium with the addition of 10 ng/ml of BDNF, CNTF and GDNF (R&D Systems) and while medium in motor neuron compared contained no neurotrophic factors to start recruitment of motor neuron axons to the muscle compartment by generation of a volumetric gradient (50 μl difference in volume between the compartments) in the device. Volumetric gradient was kept for every medium change, done every other day. Co-cultures were fixed at day 21 post-seeding for visualization of motor axon-muscle synaptic contacts.

### FACS analyses

We used an *Hb9:*:GFP reporter stem cell line previously described infected with the Ngn2 lentiviral constructs as described above (Di Giorgio et al., 2007). Briefly, cells were differentiated in 24 well plates and subjected to different patterning molecules. At each time point, cells were dissociated with Accutase (Gibco) as previously described, each replicate was frozen in Cryostor® CS10 (STEMCELL Technologies). After all samples were collected, cells were thawed in separated tubes are resuspended in sorting buffer as described by others (Klim et al., 2019), The BD FACS Aria II cell sorted was used to quantify the percentage of *Hb9*::GFP^+^ cells in each sample after using DAPI signal to determine cell viability.

### Immunofluorescence assays

Cells were washed once with PBS, fixed with 4% PFA for 20 minutes, washed again in PBS and blocked for one hour in 0.1% Triton in PBS with 10% donkey serum. Fixed cells were then washed and incubated overnight with primary antibodies at 4°C. Primary antibody solution was washed and cells were subsequently incubated with secondary antibodies (1:2000, Alexa Fluor, Life Technologies) at room temperature for 1 hour, washed with PBS and stained with DAPI. Primary antibodies used: Tuj1 (R&D, MAB1195), Islet1 (Abcam, ab178400), MAP2 (Abcam, ab5392), Synapsin (Millipore, AB1543), SMI-32 (BioLegend, 801702), Chat (Millipore, AB144P), Foxp1 (Abcam, ab16645), AnkyrinG (Millipore, MABN466). Images were analysed using FIJI.

### RNA extraction and RT-qPCR analyses

RNA was extracted with the miRNeasy Mini Kit (Qiagen, 217004). cDNA was produced with iScript kit (BioRad) using 50 ng of RNA. RT-qPCR reactions were performed in triplicates using 20 ng of cDNA with SYBR Green (BioRad) and were run on a CFX96 Touch™ PCR Machine for 39 cycles at: 95°C for 15s, 60°C for 30s, 55°C for 30s.

### Western blots

For WB analyses, cells were lysed in RIPA buffer with protease inhibitors (Roche). After protein quantification by BCA assay (ThermoFisher), ten micrograms of protein were preheated in Laemmli’s buffer (BioRad), loaded in 4-20% mini-PROTEAN® TGX™precast protein gels (BioRad) and gels were transferred to a PDVF membrane. Membranes were blocked in Odyssey Blocking Buffer (Li-Cor) and incubated overnight at 4°C with primary antibodies (1:1000 dilution). After washing with TBS-T, membranes were incubated with IRDye® secondary antibodies (Li-Cor) for one hour and imaged with Odyssey® CLx imaging system (Li-Cor). Primary antibodies used: Tuj1 (R&D, MAB1195), Synapsin (Millipore, AB1543), PSD-95 (Neuromab, 75-028), GAPDH (Millipore, MAB374).

### Multi Electrode Array analysis

Electrophysiological recordings were obtained by Axion Biosystems Multi-Electrode Array (MEA) plate system (Axion Biosystems, 12 wells or 48 wells formats) that recorded extracellular spike potential. On day 7 of differentiation, cells were detached and counted and mixed with murine glia as described above. MEA plates were previously coated with Matrigel (Corning) and cells were seeded in Neurobasal medium supplemented with ROCK inhibitor for 24 hours. Recordings were performed every 2-3 days and medium was changed after recordings. Analysis was performed with AxIS (Axion Biosystems – Neuronal Metric Tool) as described by others (Nehme et al., 2018).

### Stem cell lines, villages, single-cell RNA-sequencing, Census-seq and Dropulation

Methodology for Census-seq and Dropulation analysis are described elsewhere (Mitchell et al., 2020), we are going to provide a brief description below:

#### Human pluripotent cell lines and village generation (Mitchell et al., 2020)

Human embryonic stem cell lines used in this study were part of a collection previously described (Merkle et al., 2017). These cell lines were exome sequenced and whole genome sequenced after minimal passaging and cultured as described above. Individual stem cell lines were cultured and differentiated into liMoNes as described above. At day 6 after doxycycline induction, when cells are mostly postmitotic, cultures were dissociated with Accutase (Gibco) and resuspended in mTeSR medium with 1 μM RoCK inhibitor Y-27632. To generate balanced “villages”, cell suspensions were counted using a Scepter 2.0 Handheld Cell Counter (Millipore Sigma) with 60 μM Scepter Cell Counting Sensor (Millipore Sigma), 0.5M viable cells from each donor cell line were mixed. At this timepoint 0.5M cells were harvested for Census-seq analysis and ensure balanced representation, the rest was plated for subsequent experiments.

#### DNA isolation and library preparation (Mitchell et al., 2020) (Wells, Nemesh, Ghosh, Mitchell et al., 2021)

Every seven days, pellets were harvested from separate wells of the “liMoNes village” after dissociation with Accutase (Gibco). Pellets were lysed and DNA precipitated and DNA was used to generate libraries using TruSeq Nano DNA Library Prep Kit (Illumina), libraries were sequenced using NextSeq 500 Sequencing System (Illumina). Generated libraries were aligned to human genome using BWA, reference genome was selected to match the genomes used to generate VCF files containing the whole-genome sequenced genotypes of each donor cell line. To exclude confounding mouse DNA from glia, a multi-organism reference was used, reads competitively aligned to both genomes and only high quality (MQ≥10) were used for assignment.

#### Census-seq analysis (Mitchell et al., 2020)

The algorithms used to assign donor contribution to villages are extensively described elsewhere and their validation is outside the scope of this publication. However, briefly the aim of Census-seq algorithms is to accurately detect and precisely quantify the contribution of donors in a mixed DNA sample to monitor population dynamics over time and/or conditions. This can be achieved systematically and inexpensively by lightly sequencing genomic DNA, the algorithms attempt to determine the donors’ mixture by determining the ratio of alleles present at every SNP. The gradient-descent algorithm can then use this data to identify the donor-mix that maximizes the likelihood of any observed sequence data. Once the best ratio is identified, the algorithms compare the computed “most likely donor mix” to a VCF file that contains whole genome-sequencing data from all stem cell lines in the collection. These VCF files contain a filtered and refined matrix with alternate alleles at each variant for every donor’s genotype. Census-seq can use this data to find a vector of donor-specific contribution (to the mix) that can explain the allele counts detected at each site in the sequencing data provided. For each site, the allele frequency is inferred using the VCF reference files and its proportion of donor in the pool of DNA can then be calculated over the total counts for that specific site. The algorithms are then able to sum the proportion of each donor’s representation at every specific site and calculate total representation of each genotype, a.k.a. donor, in the pooled DNA, providing us an estimate of the ratio of donors in the village.

#### Dropulation: scRNA-sequencing and donor assignment (Wells, Nemesh, Ghosh, Mitchell et al., 2021)

For single-cell analyses, cells were harvested and prepared with 10X Chromium Single Cell 3’ Reagents V3 and sequenced on a NovaSeq 6000 (Illumina) using a S2 flow cell at 2 × 100bp. Raw sequence files were then aligned and prepared following previous Drop-seq workflow (Macosko et al., 2015). Human reads were aligned to GRCh18 and filtered for high quality mapped reads (MQ≥10) as for Census-seq standards. In order to identify donor identity of each droplet, variants were filtered through several quality controls as described previously be included in the VCF files (Wells et al., 2021), to summarise the goal is to only use sites that unambiguously and unequivocally can be detected as A/T or G/C. Once both the sequenced single-cell libraries and VCF reference files are filtered and QC’ed, the Dropulation algorithm is run. Dropulation analyses each droplet, hence a cell, independently and for each cell generates a number representing the likely provenance of each droplet from one donor. Each variant site is assigned a probability score for a given allele in the sequenced unique molecular identifier (UMI) calculated as the probability of the base observed compared to expected based, and 1 – probability that those reads disagree with the base sequenced. Donor identity is then assigned as the computed diploid likelihood at each UMI summed up across all sites.

This probability-based analysis allows to increase confidence in donor detection per barcode by increase the numbers of individuals in the VCF files: more individuals, more UMIs with site variants, more confident scores, higher quality donor assignments. After assigning a “likelihood score”, sites where only few donors have detected reads are ignored and scores are adjusted to allow only high confidence variant sites to be included. This second computer score is then added to the original likelihood as a weighted average score, this mixed coefficient defines the proportion of the population that presents each genotype and in adds to 1. Based on this mixed coefficient that takes into account reads mapped to each donors and the confidence to which each site can be used for this assignment, Dropulation then contains algorithms able to detect “doublets”, barcoded droplets with genetic DNAs assigned to two different donors, to avoid analysing barcodes with admixed identity but also to avoid excluding barcoded droplets with unclear donor assignment based on the coefficient previously calculated (Wells et al., 2021).

Once scores are calculated, the algorithm assigns donors to single droplets. Then runs the double detection and cells that are likely doublets are filtered out. After that, donor identities are confirmed only if p-value<0.05. These cells are then validated by crossing proportions of each donors as known inputs in the village and excluding any unexpected identity. Donors composing less then 0.2% of the libraries are excluded from the experiment (Wells et al., 2021).

More details on the preparation and filtering of libraries and on donor identification can be found in previously published work (Wells et al., 2021).

---

**Supplementary Figure 1.**
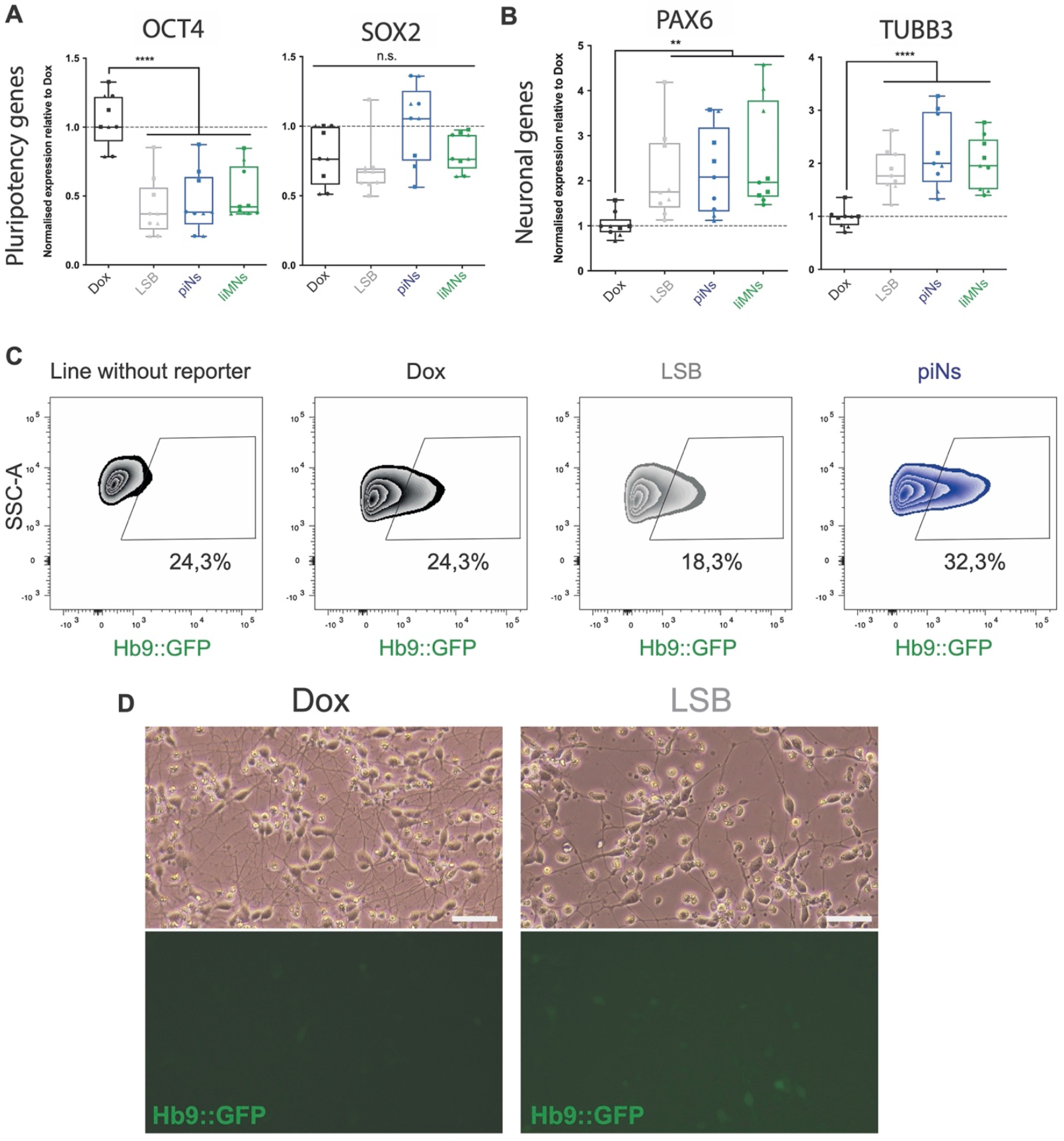
Ngn2 neuronal differentiation can be directed to different neuronal fates by small molecules patterning. (A-B) RT-qPCR quantification of pluripotency genes and genes involved in pan-neuronal development (p-values from one-way ANOVA). (C) Flow cytometry quantification of Hb9::GFP positive cells by day 4 for the other conditions. (D) Hb9::GFP expression at day7 post-induction in original Ngn2-induced Dox and LSB conditions (scale bar 50 micron).

**Supplementary Figure 2.**
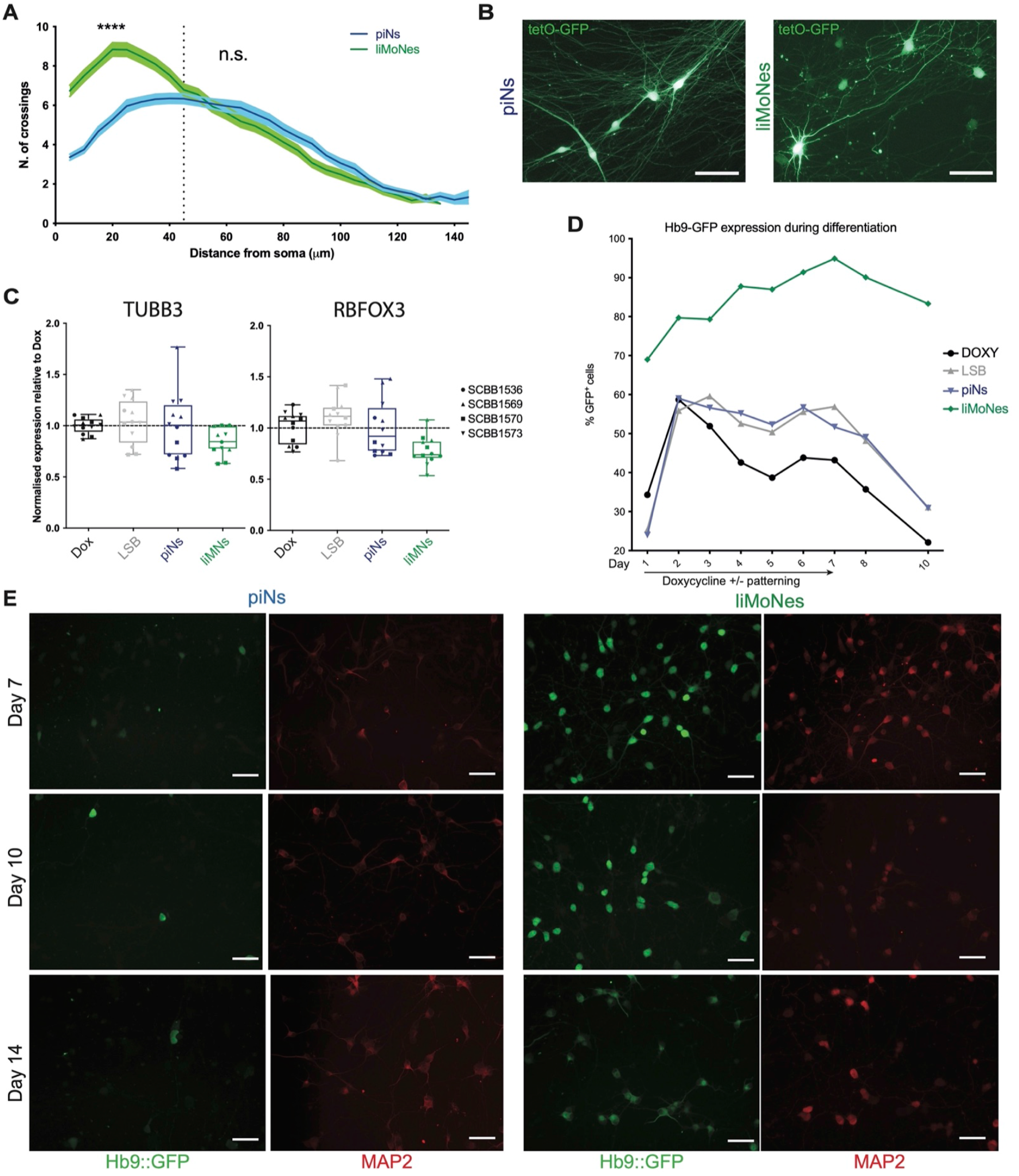
Patterned Ngn2-induced neuronal fate is maintained throughout the differentiation. (A) Quantification of arborization of piNs and liMoNes. (B) Viral tetO-GFP imaging at day30 in piNs and liMoNes, showing different cell morphology (scale bar 50 micron). (C) RT-qPCR quantification of pan-neuronal markers (p-values from one-way ANOVA). (D) Flow cytometry quantification of Hb9::GFP along differentiation. (E) Images of Hb9::GFP expression at day7, 10 and 14 post-induction in piNs and liMoNes by immunofluorescence.

**Supplementary Figure 3.**
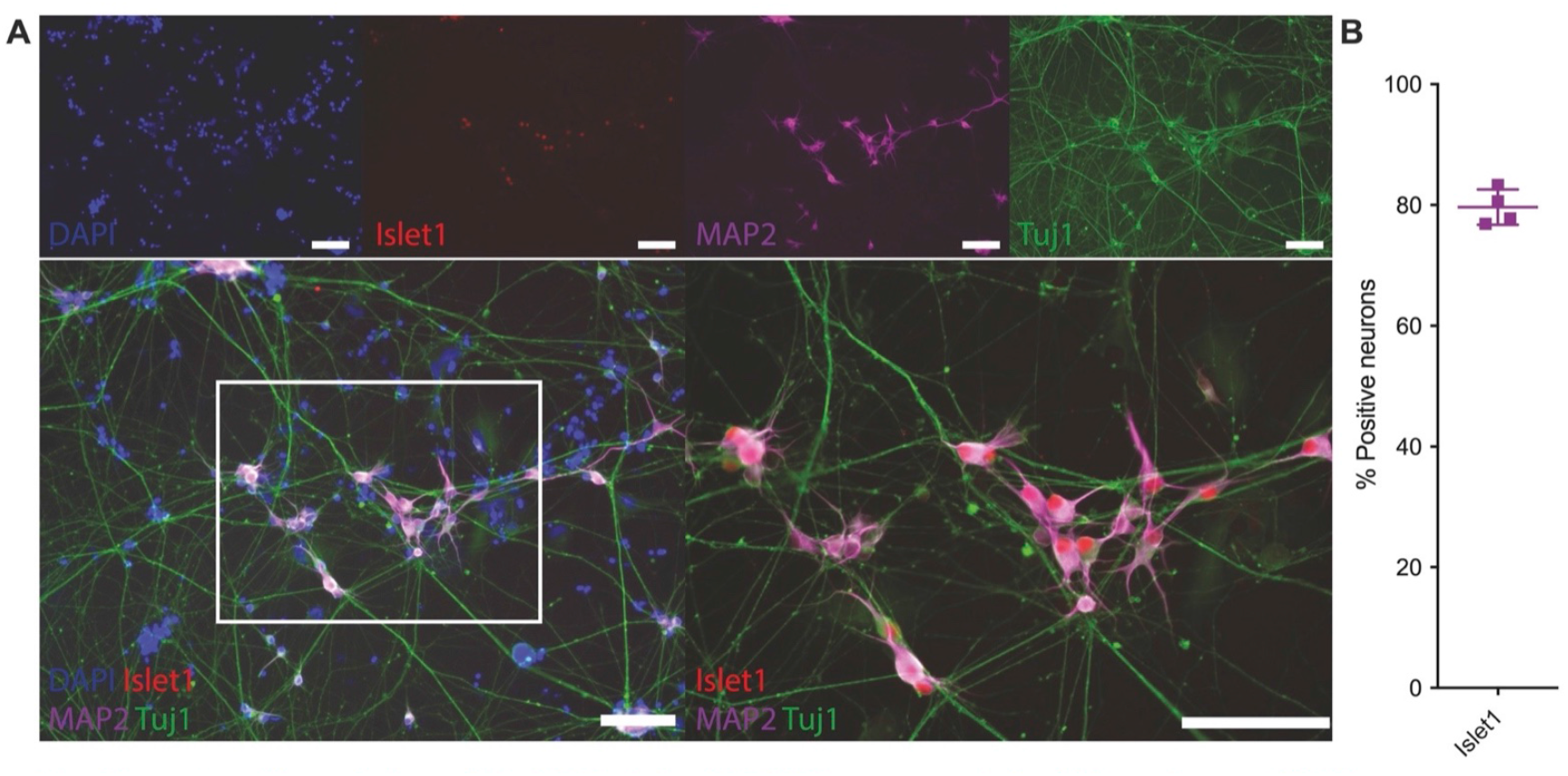
Bona fide, hPSC-derived 2D MN express similar MN markers as liMoNes. (A) lmmunofluorescence analysis for cholinergic transcription factor lslet1 and neuronal cytoskeletal proteins MAP2 and TUBB3 (tuj1) in 2D MN (scale bar 50 micron). (B) Quantification for cells in A.

**Supplementary Figure 4.**
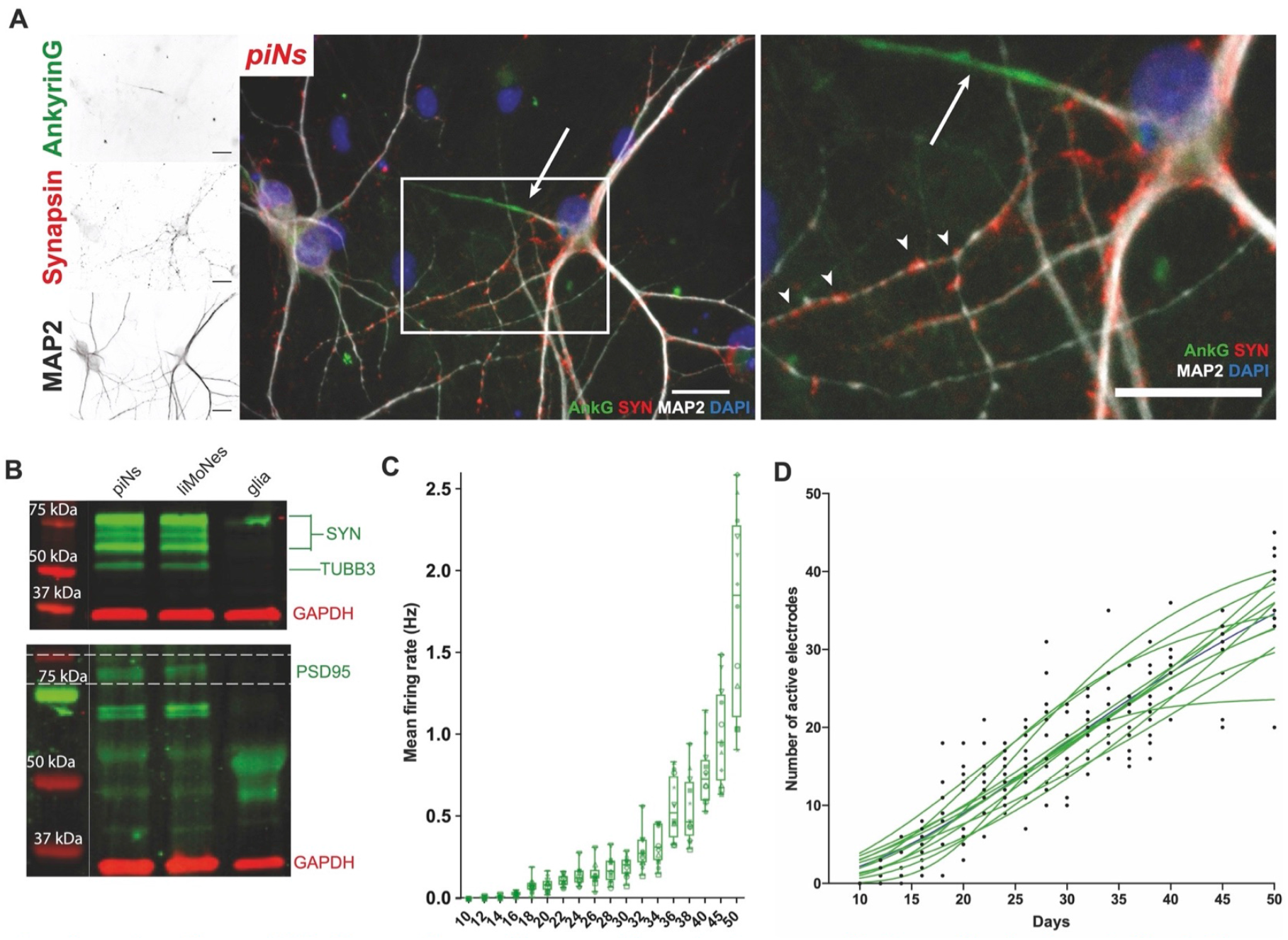
liMoNes can form active synaptic contacts comparable to previously characterised piNs. (A) lmmunofluorescence for proteins involved in the formation of functional axons and synaptic structure in piNs. (B) Western blot analysis shows expression of pre- and post-synaptic density molecules in both cell types. (C-D) Network activity of liMoNes.(C) Mean number of spikes in 10-s period in liMoNes co-cultured with murine cortical glial preparations. (D) Proportion of active electrodes detecting spontaneous activity throughout the differentiation (days). Data fit by sigmoidal function (green), median sigmoidal in black.

**Supplementary Figure 5.**
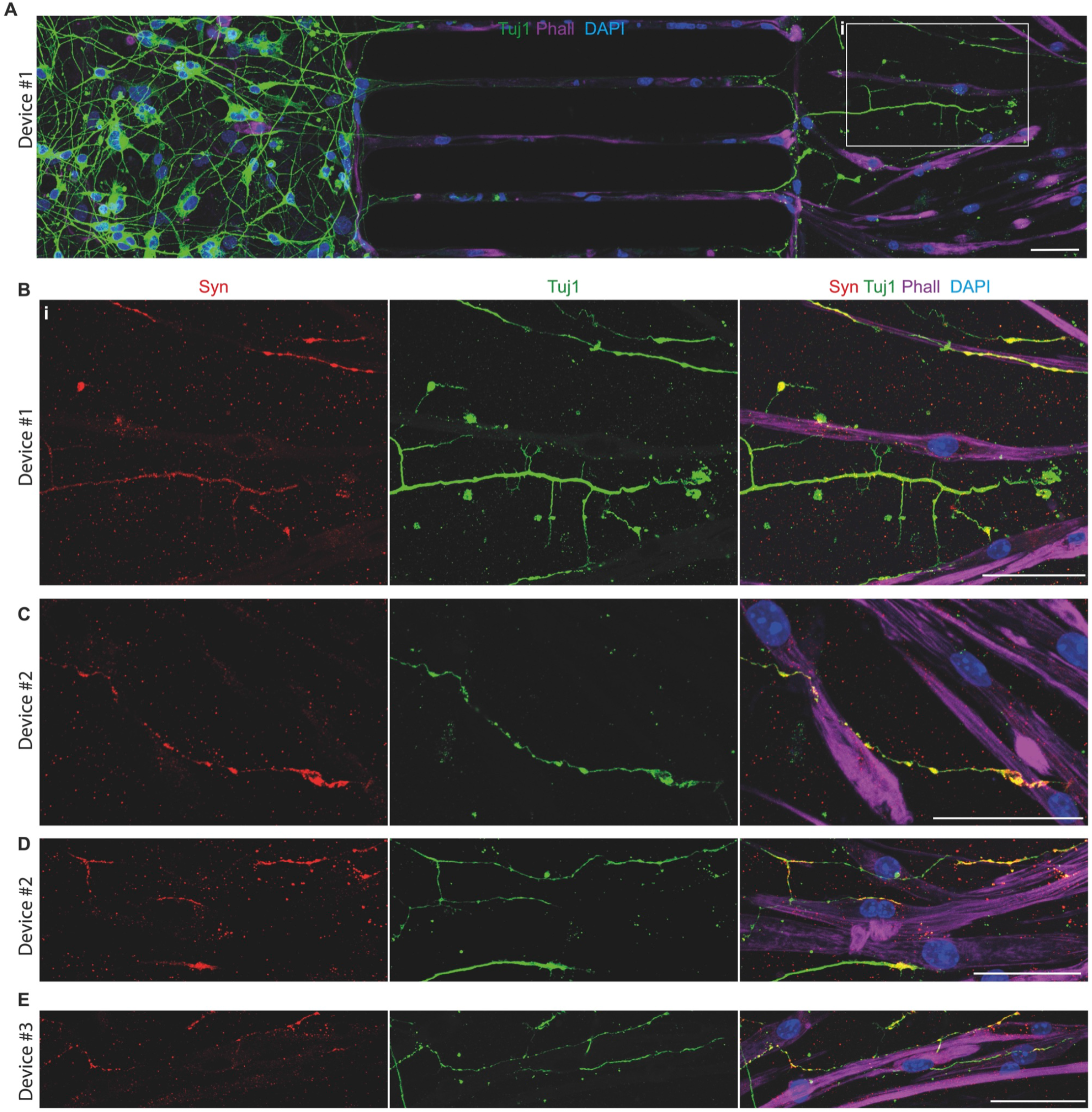
liMoNes can form NMJ-like structures *in vitro*. lmmunofluorescence of co-cultures of liMoNes and primary murine myoblasts from three devices from separate rounds of differentiation. (A) Representative image of neurons extending axons through the channels (middle), contacting primary muscle cells (rigth). (B) Insets of (A) showing liMoNes forming synaptic-like contacts with muscles cells. (C-E) Representative images from separate devices showing liMoNes forming synaptic-like contacts with muscles cells.

**Supplementary Figure 6.**
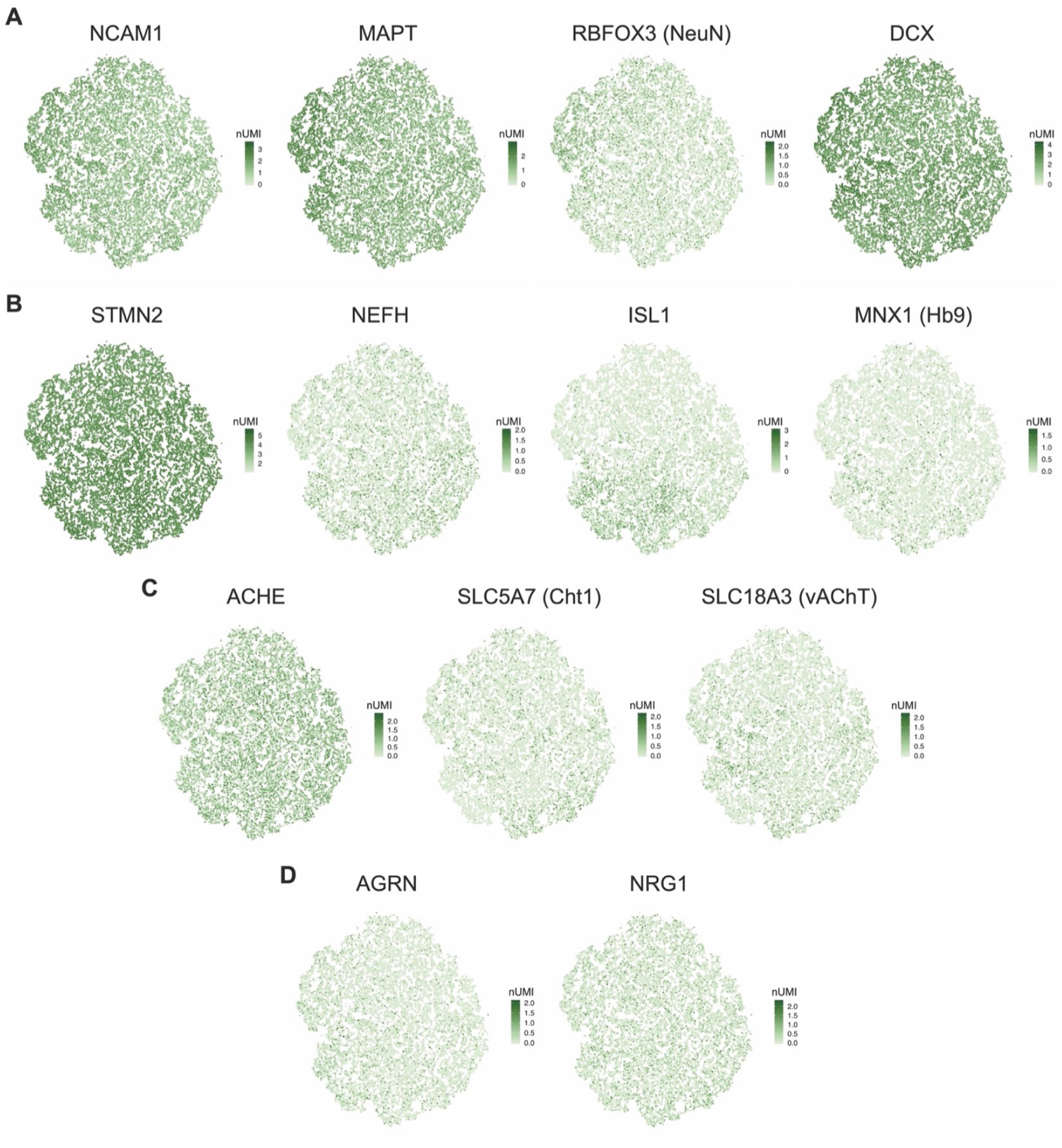
scRNAseq confirms expression of MN markers. (A) *t*-SNE projection with expression of markers specific for post-mitotic neurons. (B) *t*-SNE projection with expression of MN-specific markers. (C) *t*-SNE projection with expression of genes involved in cholinergic machinery. (D) *t*-SNE projection with expression of genes involved NMJ formation.

**Supplementary Figure 7.**
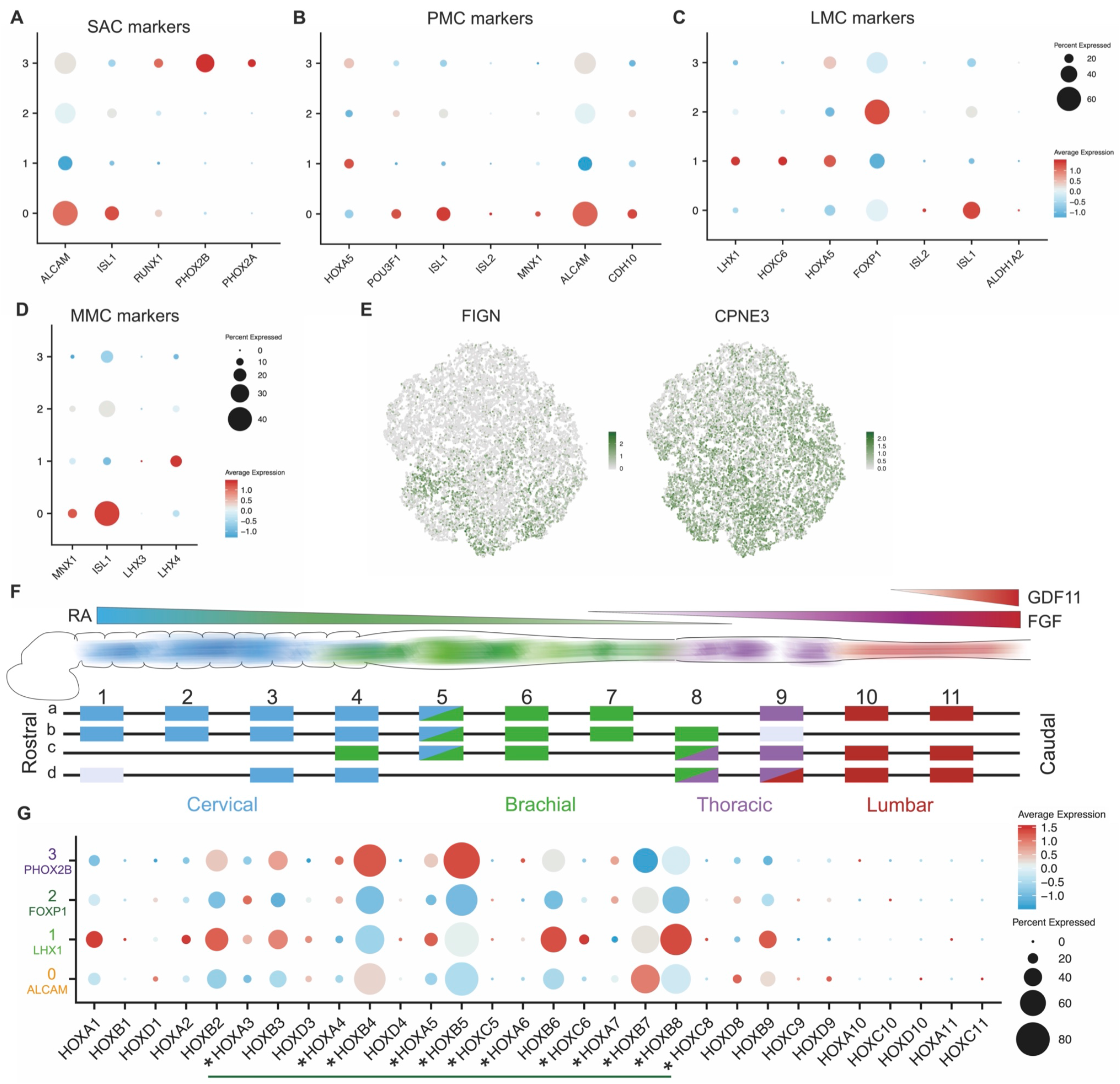
HOX genes expression and sub-columnar localization correspond to retinoid stimulation. (A) Dotplot for expression of markers specific for SAC - Spinal Accessory Column. (B) Dotplot for expression of markers specific for PMC - Phrenic Motor Column. (C) Dotplot for expression of markers specific for LMC - Lateral Motor Column. (D) Dotplot for expression of markers specific for MMC - Median Motor Column. (E) *t*-SNE projection with expression of markers expressed by digit-innervating motor neurons. (F) Schematic of spinal cord HOX genes expression. (G) Dotplot for gene expression of all HOX genes detected in the four subclusters confirms activation of retinoid depend Hox activation in vertebrates (green line) and specifically expressed in ventral spinal cord MNs (asterisks).

